# Reduced function of the glutathione S-transferase S1 suppresses behavioral hyperexcitability in *Drosophila* expressing a mutant voltage-gated sodium channel

**DOI:** 10.1101/2020.01.21.906156

**Authors:** Hung-Lin Chen, Junko Kasuya, Patrick Lansdon, Garrett Kaas, Hanxi Tang, Maggie Sodders, Toshihiro Kitamoto

## Abstract

Voltage-gated sodium (Na_v_) channels play a central role in the generation and propagation of action potentials in excitable cells such as neurons and muscles. To determine how the phenotypes of Na_v_-channel mutants are affected by other genes, we performed a forward genetic screen for dominant modifiers of the seizure-prone, gain-of-function *Drosophila melanogaster* Na_v_-channel mutant, *para^Shu^*. Our analyses using chromosome deficiencies, gene-specific RNA interference, and single-gene mutants revealed that a null allele of *glutathione S-transferase S1* (*GstS1*) dominantly suppresses *para^Shu^* phenotypes. Reduced *GstS1* function also suppressed phenotypes of other seizure-prone Na_v_-channel mutants, *para^GEFS+^* and *para^bss^*. Notably, *para^Shu^* mutants expressed 50% less *GstS1* than wild-type flies, further supporting the notion that *para^Shu^* and *GstS1* interact functionally. Introduction of a loss-of-function *GstS1* mutation into a *para^Shu^* background led to up- and down-regulation of various genes, with those encoding cytochrome P450 (CYP) enzymes most significantly over-represented in this group. Because *GstS1* is a fly ortholog of mammalian hematopoietic prostaglandin D synthase, and in mammals CYPs are involved in the oxygenation of polyunsaturated fatty acids including prostaglandins, our results raise the intriguing possibility that bioactive lipids play a role in *GstS1-*mediated suppression of *para^Shu^* phenotypes.

## INTRODUCTION

Defects in ion-channel genes lead to a variety of human disorders that are collectively referred to as channelopathies. These include cardiac arrhythmias, myotonias, forms of diabetes and an array of neurological diseases such as epilepsy, familial hyperekplexia, and chronic pain syndromes (Rajakulendran *et al*. 2012; Venetucci *et al*. 2012; Waxman and Zamponi 2014; Dib-hajj *et al*. 2015; Jen *et al*. 2016). The advent of genome-wide association studies and next-generation sequencing technology has made the identification of channelopathy mutations easier than ever before. However, the expressivity and disease severity are profoundly affected by interactions between the disease-causing genes and gene variants at other genetic loci. The significance of gene-gene interactions in channelopathies was demonstrated by Klassen et al. (2011), who performed extensive parallel exome sequencing of 237 human ion-channel genes and compared variation in the profiles between patients with the sporadic idiopathic epilepsy and unaffected individuals. The combined sequence data revealed that rare missense variants of known channelopathy genes were prevalent in both unaffected and disease groups at similar complexity. Thus, the effects of even deleterious ion-channel mutations could be compensated for by variant forms of other genes (Klassen *et al*. 2011).

*Drosophila* offers many advantages as an experimental system to elucidate the mechanisms by which genetic modifiers influence the severity of channelopathies because of the: wealth of available genomic information, advanced state of the available genetic tools, short life cycle, high fecundity, and evolutionary conservation of biological pathways (Hales *et al*. 2015; Ugur *et al*. 2016). In the current study, we focused on genes that modify phenotypes of a voltage-gated sodium (Na_v_)-channel mutant in *Drosophila*. Na_v_-channels play a central role in the generation and propagation of action potentials in excitable cells such as neurons and muscles (Hodgkin and Huxley 1952; Catterall 2012). In mammals, the Na_v_-channel gene family comprises nine paralogs. These genes encode large (∼260 kDa) pore-forming Na_v_-channel α-subunits, Na_v_1.1-Na_v_1.9, all of which have distinct channel properties and unique patterns of expression involving both subsets of neurons and other cell types. The *Drosophila* genome contains a single Na_v_-channel gene, *paralytic* (*para*), on the X chromosome. It encodes Na_v_-channel protein isoforms that share high amino-acid sequence identity/similarity with mammalian counterparts (e.g., 45%/62% with the human Na_v_ 1.1). High functional diversity of *para* Na_v_ channels is achieved through extensive alternative splicing that produces a large number (∼60) of unique transcripts (Kroll *et al*. 2015).

A number of *para* mutant alleles have been identified in *Drosophila*. They display a variety of physiological and behavioral phenotypes: lethality, olfactory defects, spontaneous tremors, neuronal hyperexcitability, resistance to insecticides, and paralysis or seizure in response to heat, cold, or mechanical shock (Suzuki *et al*. 1971; Ganetzky and Wu 1982; Lilly *et al*. 1994; Martin *et al*. 2000; Lindsay *et al*. 2008; Parker *et al*. 2011; Sun *et al*. 2012; Schutte *et al*. 2014; Kaas *et al*. 2016). One of these more recently characterized Na_v_-channel gene mutants, *para^Shu^*, is a dominant gain-of-function allele formerly referred to as *Shudderer* due to the “shuddering” or spontaneous tremors it causes (Williamson 1971; Williamson 1982). This allele contains a missense mutation that results in the replacement of an evolutionarily conserved methionine residue in Na_v_-channel homology domain III (Kaas et al. 2016). Adult *para^Shu^* mutants exhibit various dominant phenotypes in addition to shuddering, such as defective climbing behavior, increased susceptibility to electroconvulsive and heat-induced seizures, and short lifespan. They also have an abnormal down-turned wing posture and an indented thorax, both of which are thought to be caused by neuronal hyperexcitability (Williamson 1982; Kaas *et al*. 2016; Kasuya *et al*. 2019). In the current study, we carried out a forward genetic screen for dominant modifiers of *para^Shu^* and found that the phenotypes are significantly suppressed by loss-of-function mutations in the *glutathione S-transferase S1* (*GstS1*) gene. To obtain insights into the mechanisms underlying this GstS1-mediated suppression of *para^Shu^* phenotypes, we also performed RNA-sequencing analysis. This revealed changes in gene expression that are caused by reduced *GstS1* function in the *para^Shu^* background.

## MATERIALS AND METHODS

### Fly stocks and culture conditions

Flies were reared at 25°C, 65% humidity in a 12 hr light/dark cycle on a cornmeal/glucose/yeast/agar medium supplemented with the mold inhibitor methyl 4-hydroxybenzoate (0.05 %). The exact composition of the fly food used in this study was described in Kasuya et al. (2019). The *Canton-S* (*CS*) strain was used as the wild-type control. *para^Shu^*, which was originally referred to as *Shudderer* (*Shu*) (Williamson 1982) and was obtained from Mr. Rodney Williamson (Beckman Research Institute of the Hope, CA). *Drosophila* lines carrying deficiencies of interest and a UAS-*GstS1* RNAi (GD16335) were obtained from the Bloomington Stock Center (Indiana University, IN) and the Vienna *Drosophila* Resource Center (Vienna, Austria), respectively. *GstS1^M26^* was obtained from Dr. Tina Tootle (University of Iowa, IA). Genetic epilepsy with febrile seizures plus (GEFS+) and Dravet syndrome (DS) flies (*para^GEFS+^* and *para^DS^*) (Sun *et al*. 2012; Schutte *et al*. 2014) were obtained from Dr. Diane O’Dowd (University of California, Irvine, CA), and *bangsenseless* (*para^bss1^*) flies were obtained from Dr. Chun-Fang Wu (University of Iowa, IA).

### Behavioral assays

#### Reactive climbing

The reactive climbing assay was performed as previously described (Kaas *et al*. 2016), using a countercurrent apparatus originally invented by Seymour Benzer (Benzer 1967). Five to seven-day-old females (∼20) were placed into one tube (tube #0), tapped to the bottom, and allowed 15 sec to climb, at which point those that had climbed were transferred to the next tube. This process was repeated a total of 5 times. After the fifth trial, the flies in each tube (#0 ∼ #5) were counted. The climbing index (CI) was calculated using the following formula: CI = Σ(Ni x i)/(5 x ΣNi), where i and Ni represent the tube number (0-5) and the number of flies in the corresponding tube, respectively. For each genotype, at least 3 groups were tested.

#### Video-tracking locomotion analysis

Five-day-old flies were individually transferred into a plastic well (15 mm diameter x 3 mm depth) and their locomotion was recorded at 30 frames per second (fps) using a web camera at a resolution of 320 x 240 pixels for 10 minutes. The last 5 minutes of the movies were analyzed using pySolo, a multi-platform software for the analysis of sleep and locomotion in *Drosophila,* to compute the x and y coordinates of individual flies during every frame (Gilestro and Cirelli 2009). When wild-type flies are placed in a circular chamber, they spend most of their time walking along the periphery (Besson and Martin 2005), resulting in circular tracking patterns. In contrast, the uncoordinated movements caused by spontaneous tremor or jerking of *para^Shu^* mutants lead to their increased presence in the center part of the chambers. The tremor frequency was therefore indirectly assessed by determining the percentage of time that fly stayed inside a circle whose radius is 74.3% of that of the entire chamber. The distance between the fly’s position and the center of the chamber was calculated using the formula (X_i_-X_c_)^2^+(Y_i_-Y_c_)^2^<13^2^ where X_i_ and Y_i_ are the coordinates of the fly, and X_c_ and Y_c_ are the coordinates of the chamber center (13 mm is 74.3 % of the chamber radius).

#### Heat-induced seizures

Newly eclosed flies were collected in groups of 20 and aged for 3 to 5 days, after which the heat-induced seizure assay was performed as previously described (Sun *et al*. 2012). Briefly, a single fly was put into a 15 x 45 mm glass vial at room temperature (Thermo Fisher Scientific, MA) and allowed to acclimate for 2 to 10 minutes. The glass vial was then submerged in a water bath at the specified temperature for 2 minutes, during which the fly was video-taped and assessed for seizure behavior every 5 seconds. Seizure behavior was defined as loss of standing posture followed by leg shaking.

#### Bang-sensitive assay

The bang-sensitive assay was carried out following a previously described protocol (Zhang *et al*. 2002). Briefly, 10 flies were raised on conventional food for 2-3 days post-eclosion. Prior to testing, individual flies were transferred to a clean vial and acclimated for 30 minutes. Next, the vials were vortexed at maximum speed for 10 seconds, and the time to recovery was measured. Recovery was defined as the ability of flies to stand upright following paralysis. At least 5 independent bang-sensitive assays were carried out for each genotype.

#### Male mating assay

Newly eclosed *para^Shu^* males with or without one or two copies of *GstS1^M26^* (i.e., *para^Shu^*/Y; +/+, *para^Shu^*/Y; *GstS1^M26^*/+, and *para^Shu^*/Y; *GstS1^M26^*/*GstS1^M26^*) were collected. Each was placed, along with 3-5 day-old wild-type (*Canton-S*) virgin females, into a plastic tube (75 x 12 mm) containing approximately 1 ml of fly food. Tubes were kept at room temperature (∼ 22°C) for two weeks, at which point they were examined for the presence of progeny.

### Gene expression analysis

RNA was purified from one-day-old female flies using Trizol solution (Ambion, Carlsbad, CA) and an RNasy column (Qiagen, Valencia, CA). Flies of four genotypes were used: (1) +/+; +/+, (2) *para^Shu^*/+; +/+, (3) +/+; *GstS1^M26^*/+, and (4) *para^Shu^*/+; *GstS1^M26^*/+. For each genotype, RNA-sequence (RNA-seq) analysis was performed (four biological replicates) by the Iowa Institute of Human Genetics (IIHG) Genomics Division (University of Iowa, Iowa). DNase I-treated total RNA (500 ng) samples were enriched for PolyA-containing transcripts by treatment with oligo(dT) primer-coated beads. The enriched RNA pool was then fragmented, converted to cDNA, and ligated to index-containing sequence adaptors using the Illumina TruSeq Stranded mRNA Sample Preparation Kit (Cat. #RS-122-2101, Illumina, Inc., San Diego, CA). The molar concentrations of the indexed libraries were measured using the 2100 Agilent Bioanalyzer (Agilent Technologies, Santa Clara, CA) and combined equally into pools for sequencing. The concentrations of the pools were measured using the Illumina Library Quantification Kit (KAPA Biosystems, Wilmington, MA) and the samples were sequenced on the Illumina HiSeq 4000 genome sequencer using 150 bp paired-end SBS chemistry.

Sequences in FASTQ format were analyzed using the Galaxy platform (https://usegalaxy.org/). The FASTQ files were first evaluated using a quality-control tool, FastQC. The sequenced reads were filtered for those that met two conditions: minimum length >20 and quality cutoff >20. After the quality control assessments were made, the reads were mapped to Release 6 of the *Drosophila melanogaster* reference genome assembly (dm6) using the STAR tool. The number of reads per annotated gene was determined by running the featureCounts tool. The differential gene expression analyses were performed using the DESeq2 tool (Love *et al*. 2014), which uses the median of ratios method to normalize counts.

The *P*-value was adjusted (*P_adj_*) for multiple testing using the Benjamini-Hochberg procedure, which controls for the false discovery rate (FDR). For functional enrichment analysis of differentially expressed genes (DEGs), we generated a list of those for which *P_adj_*<0.05 and applied it to the GOseq tool for gene ontology analysis (Young *et al*. 2010).

### Statistical analysis

Statistical tests were performed using Sigma Plot (Systat Software, San Jose, CA). For multiple groups that exhibit non-normal distributions, the Kruskal-Wallis one-way ANOVA on ranks test was performed using Dunn’s method *post hoc*. Data that did not conform to a normal distribution are presented as box-and-whisker plots (boxplots). Values of the first, second, and third quartiles (box) are shown, as are the 10^th^ and 90^th^ percentiles (whisker), unless otherwise stated. Two-way repeated measures ANOVA and Holm-Sidak multiple comparisons were used to analyze temperature-induced behavioral phenotypes. Fisher’s exact test was used to analyze the wing and thorax phenotypes of *para^Shu^* mutants. For multiple comparison, the *P*-values were compared to the Bonferroni adjusted type I error rate for significance. Statistical analyses for RNAseq experiments are described in the previous section “Gene expression analysis by RNA-sequencing”.

## RESULTS

### The chromosomal region 53F4-53F8 contains a dominant modifier(s) of *para^Shu^*

To identify genes that interact with *para^Shu^* and influence the severity of the phenotype, we performed a forward genetic screen for dominant modifiers of *para^Shu^* using the Bloomington Deficiency Kit (Cook *et al*. 2012; Roote and Russell 2012). Females heterozygous for *para^Shu^* (*para^Shu^*/*FM7*) were crossed to males carrying a deficiency on the second or third chromosome (+/Y; *Df(2)*/balancer or +/Y;; *Df(3)*/balancer). The effects of the deficiency on *para^Shu^* were evaluated by examining the F1 female progeny trans-heterozygous for *para^Shu^* and the deficiency (e.g., *para^Shu^*/+; *Df*/+) for their reactive climbing behavior (see Materials and Methods). As reported previously, *para^Shu^* heterozygous females have a severe defect in climbing behavior due to spontaneous tremors and uncoordinated movements (Kaas *et al*. 2016). Our initial screen identified several chromosomal deficiencies that significantly improved the climbing behavior of *para^Shu^* females (Supplemental Table 1; deficiencies that resulted in CI>0.4 are shaded). The current study focuses on one of these deficiencies, *Df(2R)P803-Δ15*.

The *Df(2R)P803-Δ15* deficiency spans chromosomal region 53E-53F11 on the right arm of the second chromosome, but a lack of nucleotide level information regarding its break points made identifying the genomic region responsible for suppression of the *para^Shu^* phenotypes challenging. Therefore, we used three additional deficiencies which overlap *Df(2R)P803-Δ15* and also have molecularly defined break points (Figure 1A). Phenotypic analysis of *para^Shu^* females crossed to these deficiencies revealed that *Df(2R)Exel6065* and *Df(2R)BSC433*, but not *Df(2R)Exel6066*, had a robust suppressing effect similar to that of *Df(2R)P803-Δ15* (Figure 1B). Of the two suppressing alleles, *Df(2R)BSC433* carries the smaller deficiency; it spans genomic region 53F4 to 53F8 (Figure 1A).

**Figure 1.**
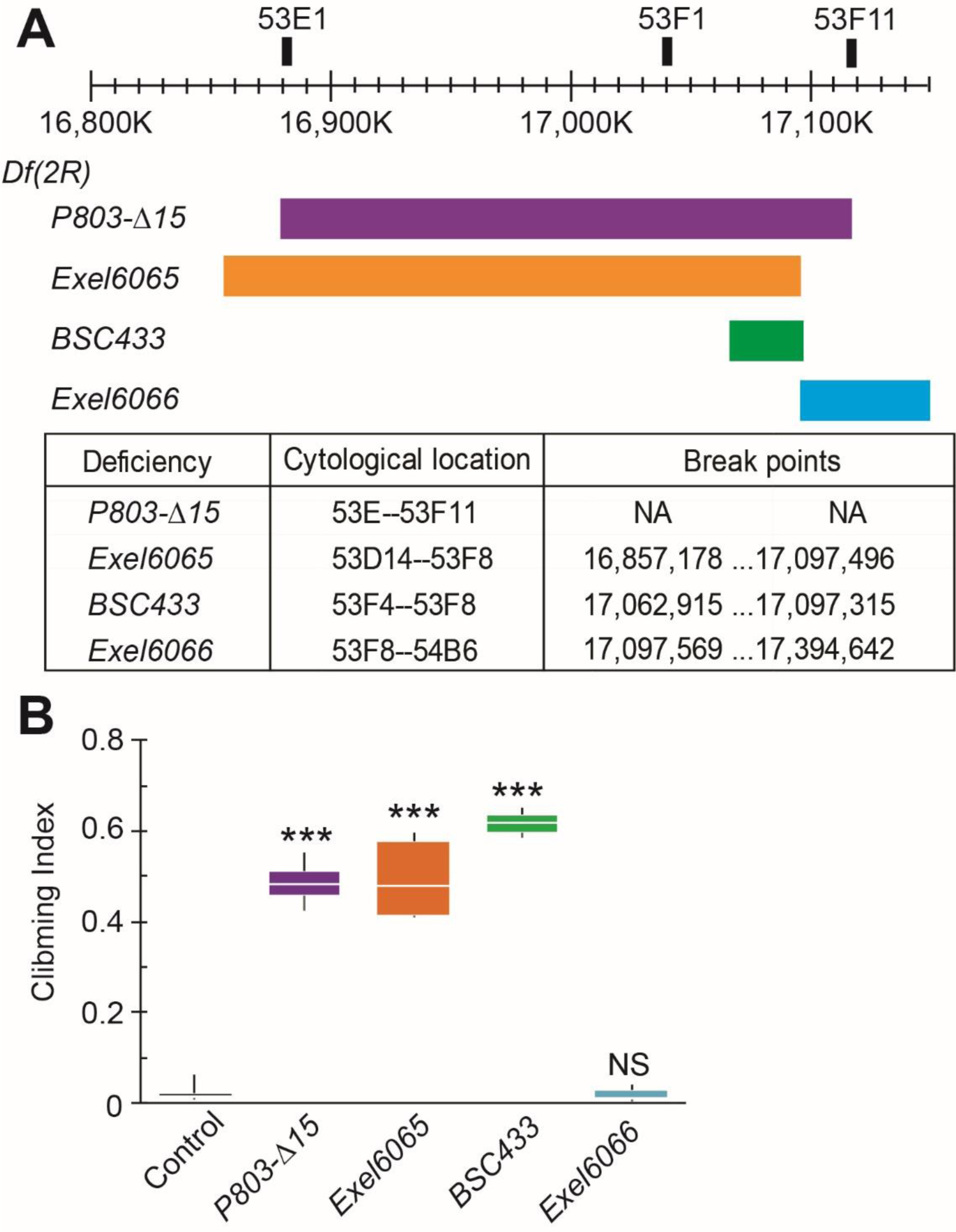
Overlapping deficiencies in the 53E-53F chromosomal region and suppression of the climbing defect of *para^Shu^* mutants. (A) Chromosomal deficiencies in 53E-53F (right arm of second chromosome) that were examined for effects on *para^Shu^* phenotypes. The cytological location and chromosomal break points of each deficiency are indicated in the table. (B) Reactive climbing behaviors of *para^Shu^* heterozygous females in the presence of the tested deficiencies. Three to eight groups of ∼20 flies per genotype were tested. The total numbers of flies tested in each group were 141 (control), 101 (*Df(2R)P803-Δ15*), 93 (*Df(2R)Exel6065*), 111 (*Df(2R)BSC433*), and 53 (*Df(2R)Exel6066*). Climbing indices are presented as box plots. The Kruskal-Wallis one-way ANOVA on ranks with Dunn’s method was used to compare between the control and deficiency groups. \*\*\**P*<0.001; NS, not significant (*P*>0.05).

The suppressive effect of *Df(2R)BSC433* was confirmed by analyzing other *para^Shu^* phenotypes. The introduction of *Df(2R)BSC433* to the *para^Shu^* background (*para^Shu^*/+; *Df(2R)BSC433*/+) significantly reduced the severity of the abnormal wing posture, indented thorax (Figure 2A), spontaneous tremors (Figure 2B), and heat-induced seizures (Figure 2C). Two deficiency lines, *Df(2R)BSC273* (49F4-50A13) and *Df(2R)BSC330* (51D3-51F9), carry a genetic background comparable to that of *Df(2R)BSC433*. Unlike *Df(2R)BSC433*, these deficiencies did not lead to suppression of *para^Shu^* phenotypes (Figure 2A-C), showing that the effect of *Df(2R)BSC433* is not due to its genetic background. Taken together, these results clearly demonstrate that removal of one copy of the genomic region 53F4-53F8 reduces the severity of multiple *para^Shu^* phenotypes, and that a dominant *para^Shu^* modifier is present in this chromosomal segment.

**Figure 2.**
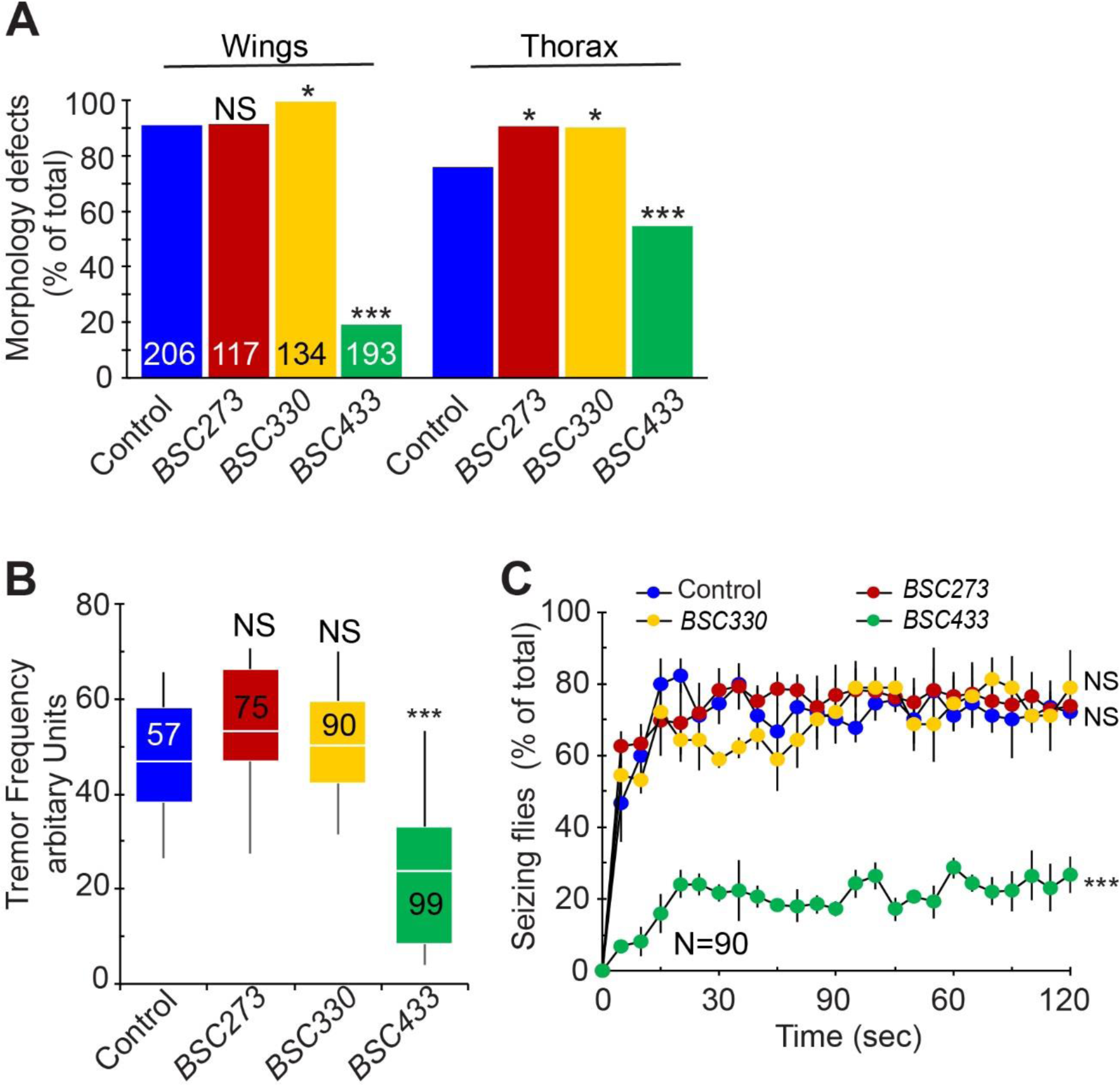
Suppression of multiple *para^Shu^* phenotypes by deletion of the genomic region 53F4-53F8. The effects of chromosomal deficiencies *Df(2R)BSC273* (49F4-50A13), *Df(2R)BSC330* (51D3-51F9), and *Df(2R)BSC433* (53F4-53F8) on *para^Shu^* phenotypes were examined. (A) Frequency of morphological defects, including down-turned wings and an indented thorax. Numbers in the bar graph indicate how many flies were scored. (B) Severity of spontaneous tremors. Numbers in the boxplot indicate how many flies were scored. (C) Severity of heat-induced seizures. Three groups of 30 flies were used per genotype. Data are shown as the averages and SEM. Fisher’s exact test with Bonferroni correction (A), the Kruskal-Wallis one-way ANOVA on ranks with Dunn’s method (B), and two-way repeated measures ANOVA and Holm-Sidak multiple comparisons (C) were used for comparisons between the control and deficiency groups. \*\*\**P*<0.001; \**P*<0.05; NS, not significant (*P*>0.05).

### *GstS1* loss of function suppresses *para^Shu^* phenotypes

Based on the molecularly defined breakpoints of *Df(2R)BSC433* (2R:17,062,915 and 2R:17,097,315), it disrupts six genes that are localized in the 53F4-53F8 region: *CG8950*, *CG6967*, *CG30460*, *CG8946* (*Sphingosine-1-phosphate lyase*; *Sply*), *CG6984*, and *CG8938* (*Glutathione S-transferase S1*; *GstS1*) (Figure 3A). To identify the gene(s) whose functional loss contributes to the marked suppression of *para^Shu^* phenotypes by *Df(2R)BSC433*, we knocked down each gene separately using gene-specific RNAi and examined the effects on *para^Shu^* phenotypes. Expression of each RNAi transgene of interest was driven by the ubiquitous Gal4 driver, *da*-*Gal4*. RNAi-mediated knockdown of *CG6967* or *Sply* resulted in developmental lethality, whereas knockdown of *CG8950*, *CG30460*, *CG6984* or *GstS1* did not. Among the viable adult progeny with gene-specific knockdown, those in which *Gst1S1* was knocked down showed the greatest improvement in wing and thorax phenotypes (Figure 3B). Thus, reduced *GstS1* function likely contributes to the suppression of *para^Shu^* phenotypes by *Df(2R)BSC433*. *GstS1^M26^* is a null allele of *GstS1* in which the entire coding region is deleted (Whitworth *et al*. 2005) and homozygotes are viable as adults. We used *GstS1^M26^* to determine how reduced *GstS1* function affects *para^Shu^* phenotypes. In *para^Shu^*/+; *GstS1^M26^*/+ flies, both the morphological (downturned wing and indented thorax) and behavioral (spontaneous tremors and heat-induced seizure) phenotypes were considerably milder than in their *para^Shu^*/+ counterparts (Figure 4A-C). *para^Shu^* phenotypes were not further improved in *GstS1^M26^* homozygotes (*para^Shu^*/+; *GstS1^M26^*/*GstS1^M26^*), where *GstS1* function was completely eliminated (Figure 4A-C). Thus, *GstS1^M26^* is a dominant suppressor of female *para^Shu^* phenotypes.

**Figure 3.**
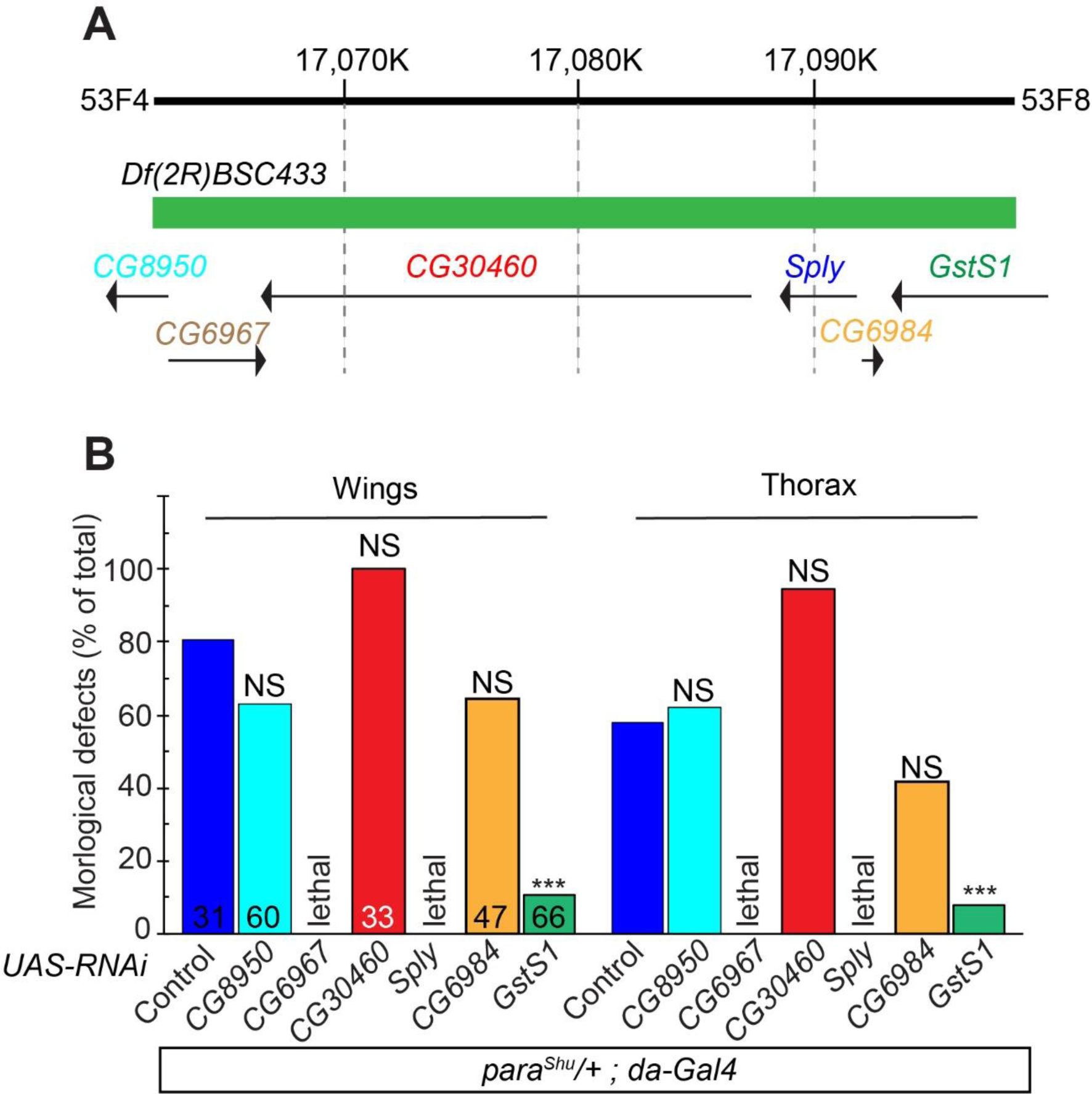
*Glutathione S-transferase S1* (*GstS1*) as a robust genetic modifier of *para^Shu^*. (A) Depiction of six genes that are localized within chromosomal region 53F4-53F8 and disrupted by the chromosomal deficiency *Df(2R)BSC433*. Arrows indicate the direction of gene transcription. (B) The frequency of *para^Shu^* morphological phenotypes following RNAi-mediated knockdown of each gene. Gene-specific RNAi was ubiquitously expressed using *da*-*GAl4* in *para^Shu^* heterozygous females (e.g., *para^Shu^*/+; *da*-*GAl4*/*UAS*-RNAi). The downturned wing (Wing) and indented thorax (Thorax) phenotypes were scored. Numbers in the bar graph indicate how many flies were scored. Fisher’s exact test with Bonferroni correction was used to analyze the data. \*\*\**P*<0.001; NS, not significant (*P*>0.05).

**Figure 4.**
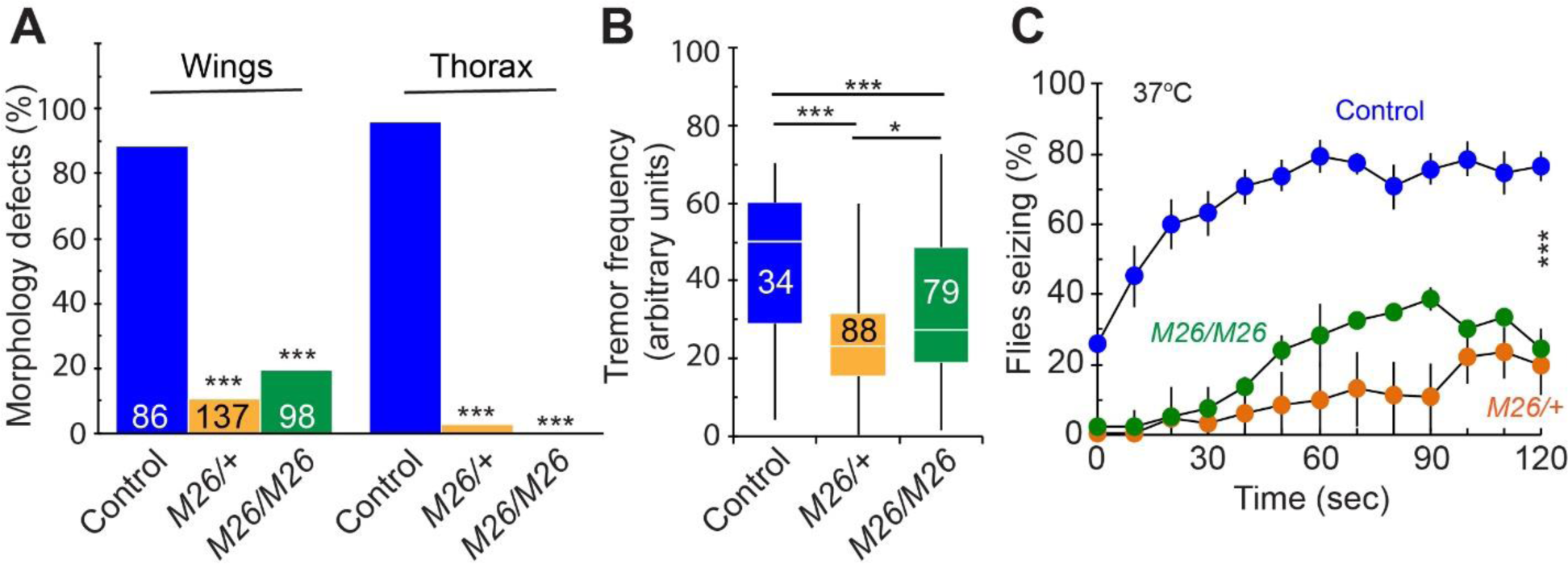
*GstS1^M26^* as a dominant suppressor of *para^Shu^* phenotypes. The effects of the *GstS1* null allele, *GstS1^M26^*, on *para^Shu^* phenotypes were examined in flies of three genotypes: (1) *para^Shu^*/+; +/+, (2) *para^Shu^*/+; *GstS1^M26^*/+, and (3) *para^Shu^*/+; *GstS1^M26^*/*GstS1^M26^*. (A) Frequencies of down-turned wings (Wings) and indented thorax (Thorax). Numbers in the bar graph indicate how many flies were scored. (B) Severity of spontaneous tremors. 8–10-day-old *para^Shu/^*+ females were used. Numbers in the boxplot indicate how many flies were scored. (C) Frequencies of heat-induced seizures. Three groups of 30-50 flies at 4-5 days after eclosion were used per genotype. Averages are shown with SEM. Fisher’s exact test with Bonferroni correction (A), the Kruskal-Wallis one-way ANOVA on ranks with Dunn’s method, (B) and two-way repeated measures ANOVA and Holm-Sidak multiple comparisons (C) were used to analyze the data. \*\*\**P*<0.001; \**P*<0.05; NS, not significant (*P*>0.05).

*GstS1^M26^* reduced the severity of the male *para^Shu^* phenotypes as well, including not only viability, but also courtship behavior and copulation. With respect to viability, *para^Shu^* males represented only 8.2% of the male progeny (*para^Shu^*/Y and *FM7*/Y) of a cross between *para^Shu^*/*FM7* females and wild-type males. Viability was significantly higher when one or two copies of *GstS1^M26^* were introduced into *para^Shu^* males (*para^Shu^*/Y; *GstS1^M26^*/+ and *para^Shu^*/Y; *GstS1^M26^*/*GstS1^M26^*), with *para^Shu^* males carrying *GstS1^M26^* representing 31.4% and 53.1% of the total male progeny, respectively (Table 1). The effects of *para^Shu^* on male courtship behavior/copulation are a consequence of the strong morphological (down-turned wings and indented thorax) and behavioral (spontaneous tremors and uncoordinated movements) phenotypes. When *para^Shu^* males were individually placed into small tubes with four wild-type virgin females and food, only one out of 43 (2.3%) produced progeny. The introduction of *GstS1^M26^* improved the ability to produce progeny; 17 out of 45 *para^Shu^* males (37.8%) heterozygous for *GstS1^M26^*, and 17 out of 44 *para^Shu^* males (38.6%) heterozygous for *GstS1^M26^*, produced progeny under the above-mentioned conditions (Table 1).

**Table 1.** Effects of *GstS1^M26^* on viability and fertility of *para^Shu^* males

| Genotype | Viability |  |  |  | Fertility |  |  |  |
| --- | --- | --- | --- | --- | --- | --- | --- | --- |
|  | Total | *FM7/Y* | *para*<sup>Shu</sup>/Y | % *para*<sup>Shu</sup>/Y | Total | Sterile | Fertile | % Fertile |
| *para*<sup>Shu</sup>/Y ; +/+ | 73 | 67 | 6 | 8.2 | 43 | 42 | 1 | 2.3 |
| *para*<sup>Shu</sup>/Y ; *GstS1*<sup>M26</sup>/+ | 121 | 83 | 38 | 31.4 | 45 | 28 | 17 | 37.8 |
| *para*<sup>Shu</sup>/Y ; *GstS1*<sup>M26</sup>/*GstS1*<sup>M26</sup> | 145 | 68 | 77 | 53.1 | 44 | 27 | 17 | 38.6 |

### Loss of function of other glutathione S-transferase genes does not suppress *para^Shu^* phenotypes as that of *GstS1*

The *Drosophila melanogaster* genome contains 36 genes that encode cytosolic glutathione S-transferases (GSTs). These are classified as Delta (D), Epsilon (E), Omega (O), Theta (T), Zeta (Z), or Sigma (S) based on similarities in the amino-acid sequences of the encoded proteins (Tu and Akgul 2005; Saisawang *et al*. 2012). *GstS1* is the sole *Drosophila* member of the S class GST genes. To determine whether reductions in the copy number of other GST genes have significant impacts on *para^Shu^* phenotypes, we generated *para^Shu^* mutants carrying chromosome deficiencies that remove the D, E, O, T, or Z class of GST genes. Given that genes encoding GSTs of the same class tend to form gene clusters, a single chromosome deficiency often removes multiple GST genes of the same class. For example, *Df(3R)Excel6164* (87B5-87B10) removes eleven GST genes of the D class (*GstD1*-*D11*) (Table 2). For GST genes on the autosomes, *para^Shu^* females (*para^Shu^*/*FM7*) were crossed to males carrying a GST deficiency on the second or third chromosome. For the two GST genes on the X chromosome (*GstT3* and *GstT4*), females carrying the deficiency (*Df*/*FM7*) were crossed to *para^Shu^* males (*para^Shu^*/Y) because males carrying this (*Df*/Y) were not viable. The female progeny carrying both *para^Shu^* and a deficiency of interest were examined for their wing posture and thorax morphology. As shown in Table 2, as well as in Figure 2, removing one copy of *GstS1* in the context of *Df(2R)BSC433* resulted in significant suppression of both the down-turned wing and the indented thorax phenotypes of *para^Shu^*, but this ability was not shared by any of the 36 other cytosolic GST genes. In some cases, however, there was partial suppression of one or the other phenotype. For example, when one copy of *GstT4* was removed (using *Df(1)Exel6245*), the wing phenotype, but not the thorax phenotype, was suppressed. Similarly, the indented thorax phenotype, but not the down-turned wing phenotype, was reduced when *GstD1-D11* was removed (using *Df(3R)Exel6164*) and when *GstT1-T2* was removed (using *Df(2R)BSC132*).

**Table 2.** Effects of GST gene deletions on wing and thorax phenotypes of *para^Shu^/+*

| Chromosomal deficiency | Deleted segment | Deleted GST genes | Flies scored | Down-turned wings |  | Indented thorax |  |
| --- | --- | --- | --- | --- | --- | --- | --- |
|  |  |  |  | (%) | (*P*-value) | (%) | (*P*-value) |
| *Df(3R)Exel6164* | 87B5-87B10 | *GstD1-D11* | 72 | 95.8 | 0.154 | 62.5 | 0.001* |
| *Df(2R)BSC335* | 55C6-55F1 | *GstE1-E11* | 59 | 91.5 | 0.741 | 86.4 | 0.550 |
| *Df(2R)BSC856* | 60E1-60E4 | *GstE12* | 68 | 77.9 | 0.208 | 83.8 | 0.397 |
| *Df(2R)BSC271* | 44F12-45A12 | *GstE13* | 67 | 94.0 | 0.480 | 77.6 | 0.0775 |
| *Df(2R)BSC273* | 49F4-50A13 | *GstE14* | 66 | 92.4 | 0.517 | 90.9 | 1 |
| *Df(3L)BSC157* | 66C12-66D6 | *GstO1-O4* | 150 | 94.7 | 0.175 | 70.0 | 0.0052 |
| *Df(2R)BSC132* | 45F6-46B4 | *GstT1-T2* | 51 | 68.6 | 0.025 | 9.80 | <0.00001* |
| *Df(1)Exel6254* | 19C4-19D1 | *GstT3* | 34 | 91.2 | 1 | 94.2 | 0.691 |
| *Df(1)Exel6245* | 11E11-11F4 | *GstT4* | 26 | 23.1 | <0.00001* | 80.8 | 0.277 |
| *Df(3R)by10* | 85D8-85E13 | *GstZ1-Z2* | 57 | 89.5 | 1 | 91.2 | 1 |
| *Df(2R)BSC433* | 53F4-53F8 | *GstS1* | 56 | 12.5 | <0.00001* | 35.7 | <0.00001* |
| No deficiency | NA | NA | 44 | 88.6 | NA | 90.9 | NA |

Statistical significance in the severity of wing and thorax phenotypes between *para*<sup>Shu</sup> (*para*<sup>Shu/+</sup>) and *para*<sup>Shu</sup> with a deficiency (*para*<sup>Shu/+</sup> ; *Df/+* or *para*<sup>Shu/Df) was assessed using Fisher's exact test. The *P*-values were compared to Bonferroni adjusted type I error rate of 0.05/11 (=0.004545.....) for significance (\*). NA, not applicable.</sup>

### *GstS1^M26^* suppresses the phenotypes of other *para* gain-of-function mutants

We next examined whether phenotypes of other Na_v_-channel mutants are similarly affected by reduced *GstS1* function. Generalized epilepsy with febrile seizures plus (GEFS+) and Dravet syndrome (DS) are common childhood-onset genetic epileptic encephalopathies (Claes *et al*. 2001; Catterall *et al*. 2010). Sun et al. (2012) and Schutte et al. (2014) created *Drosophila para* knock-in alleles, gain-of-function *para^GEFS+^* and loss-of-function *para^DS^*, by introducing a disease-causing human GEFS+ or DS mutation at the corresponding position of the fly Na_v_-channel gene. At 40°C, *para^GEFS+^* homozygous females and hemizygous males exhibit a temperature-induced seizure-like behavior that is similar to, but milder than, that observed in *para^Shu^* flies (Sun *et al*. 2012; Kaas *et al*. 2016; Kasuya *et al*. 2019). *para^DS^* flies lose their posture shortly after being transferred to 37°C (Schutte *et al*. 2014). The temperature-induced phenotype of *para^GEFS+^* was significantly suppressed when a single copy of *GstS1^M26^* was introduced into *para^GEFS+^* males (*para^GEFS+^*/Y; *GstS1^M26^*/+) (Figure 5A). In contrast, the severity of the phenotype in *para^DS^* males was unaffected by a copy of *GstS1^M26^* (*para^GEFS+^*/Y; *GstS1^M26^*/+) (Figure 5B).

**Figure 5.**
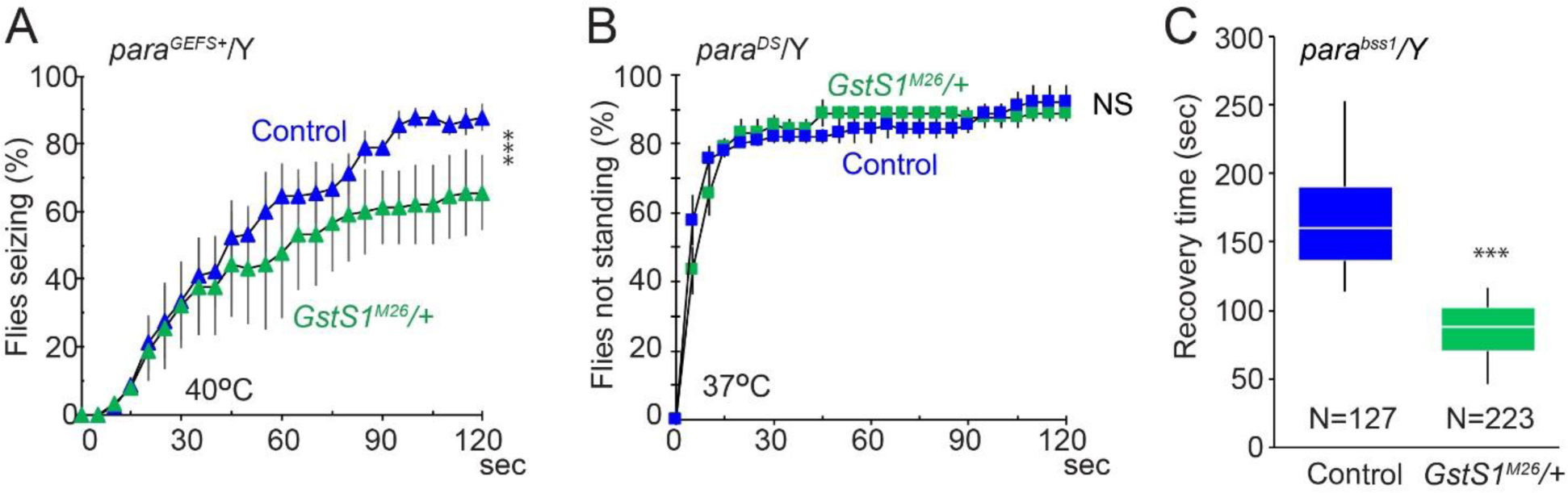
Phenotypes of other neurological mutants are suppressed by *GstS1^M26^*. (A) Frequencies of heat-induced seizure at 40°C in *para^GEFS+^* plus *GstS1^M26^* males (*para^GEFS+^*/Y; *GstS1^M26^*/+) or *para^GEFS+^* males (*para^GEFS+^*/Y; +/+). (B) Frequencies of *para^DS^* males that did not stand at 37°C. For (A) and (B), averages of 3 experiments and SEM are shown. In each experiment, 30 flies were examined. (C) Recovery time required for *para^bss1^* plus *GstS1^M26^* males (*para^bss1^*/Y; *GstS1^M26^*/+) and *para^bss1^* males (*para^bss1^*/Y; +/+) to recover from paralysis induced by mechanical shock. Data are presented as box plots. Total numbers of flies observed were 127 and 223 flies for *para^bss1^*/Y; +/+ and *para^bss1^*/Y; *GstS1^M26^*/+, respectively. Data analysis involved two-way repeated measures ANOVA and Holm-Sidak multiple comparisons (A and B) and the Mann-Whitney *U* test (C). \*\*\**P*<0.001; \**P*<0.05; NS, not significant (*P*>0.05).

We also examined *para^bss1^*, which is a hyperexcitable, gain-of-function *para* mutant allele that displays semi-dominant, bang-sensitive paralysis (Parker *et al*. 2011). The severity of the *para^bss1^* bang-sensitivity was evaluated as the time for recovery from paralysis that had been induced by mechanical stimulation (10 seconds of vortexing). All *para^bss1^* flies were paralyzed immediately after this mechanical stimulation. By three minutes after mechanical stimulation, 92% of the *para^bss1^* males carrying *GstS1^M26^* (*para^bss1^*/Y; *GstS1^M26^*/+) had recovered from paralysis and were able to right themselves, whereas only 12.6% of *para^bss1^* males had recovered. The median recovery time for *para^bss1^* males carrying *GstS1^M26^* was 88 seconds and that for *para^bss1^* males was 160 seconds (Figure 5C).

### RNA sequencing analysis revealed changes in gene expression caused by *para^Shu^* and *GstS1^M26^* mutations

To gain insights into the molecular basis of the *GstS1*-dependent suppression of *para^Shu^* phenotypes, we performed RNA sequencing (RNA-seq) analysis and examined the transcriptome profiles of *para^Shu^* and wild-type females with or without *GstS1^M26^*. Whole-body transcriptomes of one-day-old females were compared among four genotypes: (1) +/+; +/+, (2) *para^Shu^*/+; +/+, (3) +/+; *GstS1^M26^/+*, and (4) *para^Shu^*/+; *GstS1^M26^*/+. Each sample generated at least 21 million sequencing reads, of which >99% met the criteria of having a quality score of >20 and a length of >20 bp. Moreover, duplicate reads encompassed ∼70% of total reads, which was expected from the RNA-seq data (Bansal 2017).

We found that 129 genes were differentially expressed (threshold: adjusted *P*-value (*P_adj_*)<0.05) between *para^Shu^* and wild-type females. Among these, 89 and 40 genes were up- and down-regulated, respectively, in *para^Shu^* vs. wild-type flies (Supplemental Table 2). Gene ontology analysis of the differentially expressed genes was performed using GOseq tools (Young *et al*. 2010). Genes associated with four Gene Ontology categories were found to be overrepresented within the dataset (*P_adj_* <0.05), each with a functional connection to the chitin-based cuticle: “structural constituent of chitin-based larval cuticle (GO:0008010)”, “structural constituent of chitin-based cuticle (GO:0005214)”, “structural constituent of cuticle (GO:0042302)”, and “chitin-based cuticle development (GO:0040003)” (Table 3A). Within these GO categories, eight genes were differentially expressed between *para^Shu^* and wild-type flies (Table 3B).

**Table 3A.** Enriched GO terms that are overrepresented in differentially expressed genes in *ara^Shu^*/+ compared with control

| Gene ontology | Term | Ontology class | *P*<sub>adj</sub> over-represented value | # of genes |
| --- | --- | --- | --- | --- |
| GO:0008010 | structural constituent of chitin-based larval cuticle | MF | 8.83E-3 | 8 |
| GO:0005214 | structural constituent of chitin-based cuticle | MF | 1.05E-2 | 8 |
| GO:0042302 | structural constituent of cuticle | MF | 1.24E-2 | 8 |
| GO:0040003 | chitin-based cuticle development | BP | 2.01E-2 | 9 |

MF: molecular function, BP: biological process.

**Table 3B.** Differentially expressed genes in *para^Shu^* /+ compared with control that are included n the enriched GO terms

| Flybase ID | Gene symbol | Fold change (log2) | Fold change | *P*<sub>adj</sub> | Gene product |
| --- | --- | --- | --- | --- | --- |
| FBgn0033603 | *Cpr47Ef* | 1.14 | 2.20 | 2.03E-11 | Cuticular protein 47Ef |
| FBgn0004782 | *Ccp84Ab* | 1.02 | 2.03 | 1.83E-10 | Ccp84Ab |
| FBgn0004783 | *Ccp84Aa* | 0.92 | 1.89 | 9.93E-05 | Ccp84Aa |
| FBgn0001112 | *Gld* | 0.73 | 1.66 | 3.22E-03 | Glucose dehydrogenase |
| FBgn0004780 | *Ccp84Ad* | 0.72 | 1.65 | 1.02E-02 | Ccp84Ad |
| FBgn0035281 | *Cpr62Bc* | 0.64 | 1.56 | 3.59E-03 | Cuticular protein 62Bc |
| FBgn0036619 | *Cpr72Ec* | 0.64 | 1.55 | 3.97E-02 | Cuticular protein 72Ec |
| FBgn0036680 | *Cpr73D* | 0.514 | 1.43 | 3.35E-02 | Cuticular protein 73D |
| FBgn0052029 | *Cpr66D* | -0.68 | 0.62 | 9.58E-03 | Cuticular protein 66D |

Among the genes that are differentially regulated (*P_adj_*<0.05) between wild-type and *para^Shu^* flies (Supplemental Table 2), 16 displayed a fold change of >2 and all are up-regulated in *para^Shu^* flies (Table 4). They encode: a transferase (*CG32581*), two lysozymes (*LysC* and *LysD*), two endopeptidases (*Jon25Bi* and *CG32523*), one endonuclease (*CG3819*), two cytochrome P450 proteins (*Cyp4p1 and Cyp6w1*), three ABC transporters (*l(2)03659, CG7300 and CG1494*), three transcription factors (*lmd, CG18446 and Ada1-1*), and two cuticle proteins (*Cpr47Ef and Ccp84Ab*). Of note, *GstS1* was one of the 40 genes that are significantly down-regulated in *para^Shu^* females; the average normalized sequence counts (DESeq2) were 50% reduced (15562.21 vs 7782.01, adjusted *P_adj_*=0.00036) (Table 5, Figure 6). In general, we did not observe any significant differences in the expression of other GST genes between *para^Shu^* and wild-type flies, with the only exceptions being *GstD2* and *GstO2* (Table 5), down-regulated and up-regulated, respectively.

**Figure 6.**
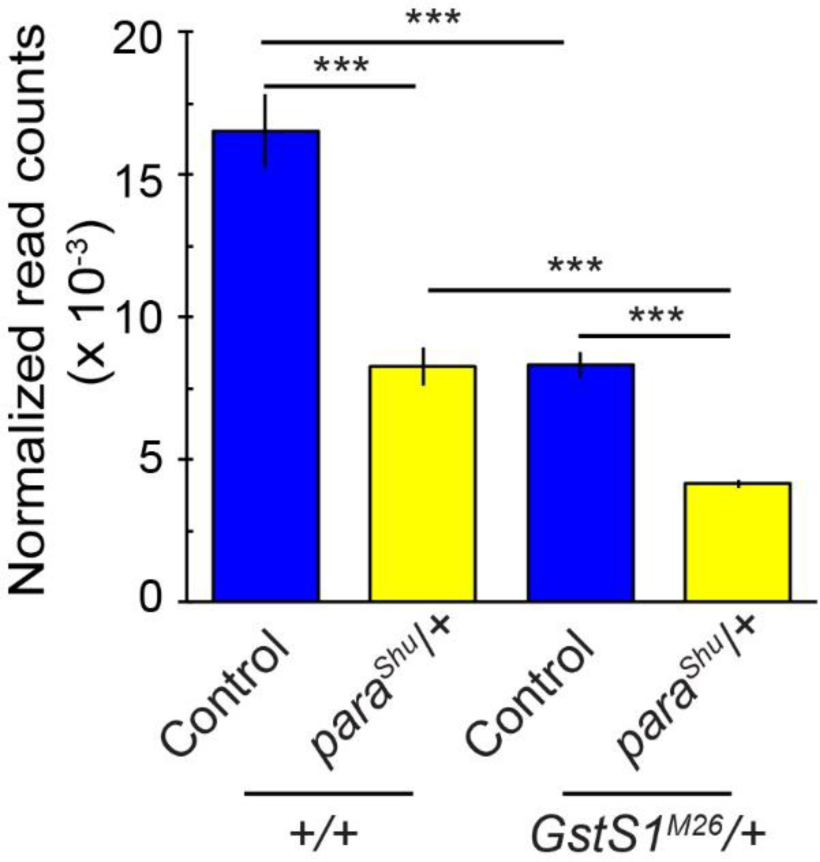
Reduction of *GstS1* expression in *para^Shu^*. Levels of *GstS1* transcript, as evaluated by RNAseq analysis in control (*Canton-S*) and *para^Shu^* heterozygous females with or without a *GstS1^M26^* mutation (*para^Shu^*/+; +/+ or *para^Shu^*/+; *GstS1^M26^*/+) (see Materials and Methods). Averages of four biological replicates are shown, as normalized read counts with SEM and adjusted *P*-values (*P_adj_*). \*\*\**P_adj_*<0.001.

**Table 4.**
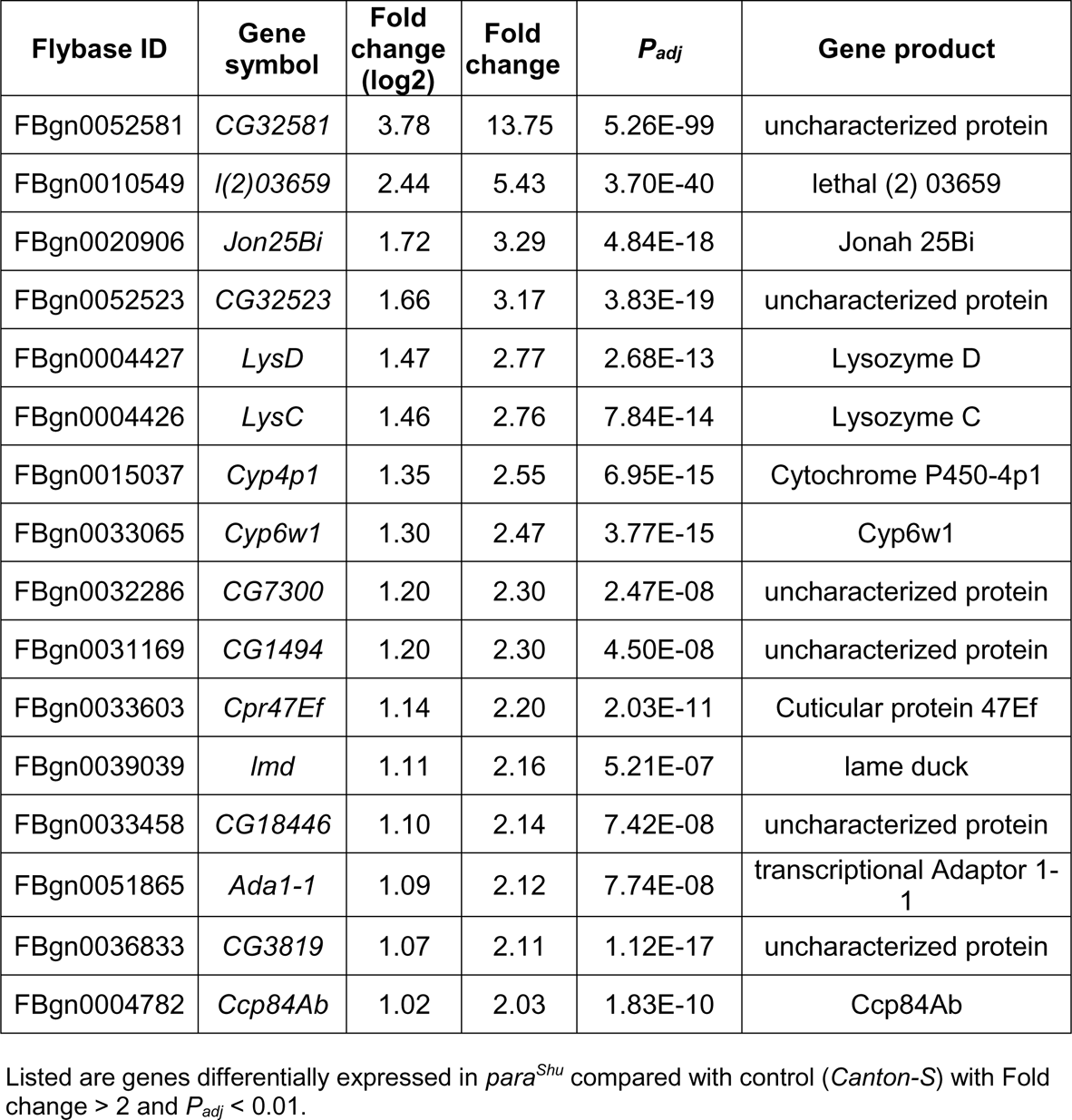
Genes most differentially expressed in *para^Shu^*/+ compared with control

**Table 5.** Expression levels of GST genes in control and *para^Shu^/+*

| GST genes | Flybase ID | average of normalized counts (DESeq2) |  | Fold change (log2) | Fold change | *P*<sub>adj</sub> |
| --- | --- | --- | --- | --- | --- | --- |
|  |  | Control | *para*<sup>Shu</sup> |  |  |  |
| *GstS1* | FBgn0063499 | 15562.21 | 7782.01 | -0.76 | 0.59 | <0.001*** |
| *GstD1* | FBgn0063495 | 10428.37 | 10078.77 | -0.02 | 0.99 | 1.000 |
| *GstD2* | FBgn0010041 | 175.19 | 54.34 | -0.72 | 0.61 | 0.010* |
| *GstD3* | FBgn0037696 | 315.48 | 208.52 | -0.20 | 0.87 | 1.000 |
| *GstD4* | FBgn0063492 | 6.43 | 5.55 | -0.06 | 0.96 | 1.000 |
| *GstD5* | FBgn0063498 | 43.77 | 20.34 | -0.32 | 0.80 | 0.878 |
| *GstD6* | FBgn0010043 | 2.45 | 4.40 | 0.14 | 1.10 | 1.000 |
| *GstD7* | FBgn0050000 | 75.87 | 83.37 | 0.07 | 1.05 | 1.000 |
| *GstD8* | FBgn0086348 | 50.70 | 55.24 | 0.06 | 1.04 | 1.000 |
| *GstD9* | FBgn0010044 | 373.48 | 431.20 | 0.10 | 1.07 | 1.000 |
| *GstD10* | FBgn0063497 | 126.21 | 176.08 | 0.30 | 1.23 | 1.000 |
| *GstD11* | FBgn0037697 | 34.52 | 43.76 | 0.18 | 1.13 | 1.000 |
| *GstE1* | FBgn0033381 | 934.78 | 894.99 | -0.05 | 0.97 | 1.000 |
| *GstE2* | FBgn0010226 | 63.43 | 77.50 | 0.18 | 1.13 | 1.000 |
| *GstE3* | FBgn0010042 | 679.13 | 697.94 | 0.02 | 1.01 | 1.000 |
| *GstE4* | FBgn0033817 | 89.15 | 101.79 | 0.14 | 1.10 | 1.000 |
| *GstE5* | FBgn0063491 | 203.55 | 227.27 | 0.10 | 1.07 | 1.000 |
| *GstE6* | FBgn0038029 | 2026.31 | 1985.35 | -0.02 | 0.99 | 1.000 |
| *GstE7* | FBgn0001149 | 561.78 | 493.62 | -0.09 | 0.94 | 1.000 |
| *GstE8* | FBgn0035906 | 410.02 | 375.98 | -0.06 | 0.96 | 1.000 |
| *GstE9* | FBgn0010039 | 887.16 | 1218.13 | 0.39 | 1.31 | 0.206 |
| *GstE10* | FBgn0063493 | 90.21 | 111.71 | 0.17 | 1.13 | 1.000 |
| *GstE11* | FBgn0042206 | 287.81 | 283.10 | -0.02 | 0.99 | 1.000 |
| *GstE12* | FBgn0030484 | 4456.11 | 4955.73 | 0.11 | 1.08 | 1.000 |
| *GstE13* | FBgn0038020 | 922.06 | 894.20 | -0.02 | 0.99 | 1.000 |
| *GstE14* | FBgn0035904 | 118.30 | 136.79 | 0.12 | 1.09 | 1.000 |
| *GstO1* | FBgn0035907 | 604.75 | 547.12 | -0.09 | 0.94 | 1.000 |
| *GstO2* | FBgn0063494 | 1408.05 | 2117.33 | 0.52 | 1.43 | 0.002** |
| *GstO3* | FBgn0034354 | 816.42 | 835.42 | 0.03 | 1.02 | 1.000 |
| *GstO4* | FBgn0050005 | 171.63 | 192.39 | 0.13 | 1.09 | 1.000 |
| *GstT1* | FBgn0031117 | 1085.64 | 998.88 | -0.07 | 0.95 | 1.000 |
| *GstT2* | FBgn0034335 | 511.60 | 386.89 | -0.32 | 0.80 | 0.705 |
| *GstT3* | FBgn0063496 | 639.30 | 587.15 | -0.11 | 0.93 | 1.000 |
| *GstT4* | FBgn0010040 | 2218.35 | 2405.08 | 0.10 | 1.07 | 1.000 |
| *GstZ1* | FBgn0027590 | 306.76 | 252.35 | -0.24 | 0.85 | 1.000 |
| *GstZ2* | FBgn0010038 | 272.48 | 246.37 | -0.09 | 0.94 | 1.000 |

Transcript levels of the 36 genes encoding soluble GSTs were evaluated by DEseq2 analysis of four biological replicates in control (*Canton-S*) and *para*<sup>*Shu*</sup>. Adjusted P-values ( $P_{adj}$ ) were obtained using Benjamini-Hochberg (BH) procedure ( $*P_{adj} < 0.05$ ; $**P_{adj} < 0.01$ ; $***P_{adj} < 0.001$ ).

We next examined how *GstS1^M26^* affects gene expression profiles in *para^Shu^* mutants. The fact that *GstS1^M26^* is a deletion mutation that removes the entire coding region of *GstS1* (Whitworth *et al*. 2005) is consistent with our discovery that the levels of the *GstS1* transcript were 50% lower than those in wild-type flies when one copy of *GstS1^M26^* was introduced (Figure 6). Since *para^Shu^* and *GstS1^M26^* each reduced *GstS1* expression by ∼50%, the level of *GstS1* expression in *para^Shu^*; *GstS1^M26^* double heterozygotes (*para^Shu^*/+; *GstS1^M26^*/+) was approximately one quarter of that in wild-type flies (Figure 6).

Comparison of *para^Shu^* flies to *para^Shu^* and *GstS1^M26^* double mutants (*para^Shu^*/+; +/+ vs *para^Shu^*/+; *GstS1^M26^*/+) revealed the differential expression of 220 genes (for *P_adj_*<0.05; Supplemental Table 2). Among these, 120 were up-regulated and 100 were downregulated in *para^Shu^* plus *GstS1^M26^* flies. Functional enrichment analysis of the differentially expressed genes revealed that genes associated with five specific molecular functions were over-represented. These include “heme binding” (GO:0020037), “tetrapyrrole binding” (GO:0046906), “iron ion binding” (GO:0005506), “oxidoreductase activity, acting on paired donors, with incorporation or reduction of molecular oxygen” (GO:0016705), and “cofactor binding” (GO:0048037) (Table 6A). Thirteen differentially regulated genes were associated with all five GO terms. These all encode heme-containing enzymes CYPs (Table 6B, marked with asterisks) that catalyze a diverse range of reactions and are critical for normal developmental processes and the detoxification of xenobiotic compounds (Hannemann et al. 2007; Isin and Guengerich 2007; Chung et al. 2009).

**Table 6A.** Enriched GO terms that are overrepresented in differentially expressed genes in *ara^Shu^/+; GstS1^M26^/+ compared with para^Shu^/+; +/+*

| Gene ontology | Term | Ontology class | *P*<sub>adj</sub> over-represented value | # of genes |
| --- | --- | --- | --- | --- |
| GO:0020037 | heme binding | MF | 7.31E-4 | 14 |
| GO:0046906 | tetrapyrrole binding | MF | 7.31E-4 | 14 |
| GO:0005506 | iron ion binding | MF | 1.53E-3 | 14 |
| GO:0016705 | oxidoreductase activity, acting on paired donors, with incorporation or reduction of molecular oxygen | MF | 2.83E-3 | 14 |
| GO:0048037 | cofactor binding | MF | 4.60E-3 | 21 |

MF: molecular function

**Table 6B.** Differentially expressed genes in *para^Shu^/+; GstS1^M26^/+ compared with para^Shu^/+; +/+* hat are included in the enriched GO terms

| Flybase ID | Gene symbol | Fold change (log2) | Fold change | *P*<sub>adj</sub> | Gene product |
| --- | --- | --- | --- | --- | --- |
| FBgn0013772 | *Cyp6a8* * | 1.69 | 3.22 | 9.37E-20 | Cytochrome P450-6a8 |
| FBgn0000473 | *Cyp6a2* * | 1.51 | 2.84 | 6.10E-15 | Cytochrome P450-6a2 |
| FBgn0041337 | *Cyp309a2* * | 1.11 | 2.16 | 1.07E-07 | Cyp309a2 |
| FBgn0033978 | *Cyp6a23* * | 1.05 | 2.07 | 7.16E-15 | Cyp6a23 |
| FBgn0033980 | *Cyp6a20* * | 0.80 | 1.74 | 4.29E-04 | Cyp6a20 |
| FBgn0033983 | *ADPS* | 0.78 | 1.72 | 9.71E-08 | Alkyldihydroxyacetone-phosphate synthase |
| FBgn0015039 | *Cyp9b2* * | 0.68 | 1.60 | 5.52E-07 | Cytochrome P450-9b2 |
| FBgn0031689 | *Cyp28d1* * | 0.66 | 1.58 | 8.28E-06 | Cyp28d1 |
| FBgn0015037 | *Cyp4p1* * | 0.59 | 1.51 | 2.79E-04 | Cytochrome P450-4p1 |
| FBgn0036381 | *CG8745* | -0.37 | 0.78 | 4.09E-02 | uncharacterized protein |
| FBgn0003965 | *v* | -0.38 | 0.77 | 4.06E-02 | vermilion |
| FBgn0035906 | *GstO2* | -0.41 | 0.75 | 1.59E-02 | Glutathione S transferase O2 |
| FBgn0036927 | *Gabat* | -0.49 | 0.71 | 1.13E-04 | gamma-aminobutyric acid transaminase |
| FBgn0000566 | *Eip55E* | -0.54 | 0.69 | 1.27E-04 | Ecdysone-induced protein 55E |
| FBgn0051674 | *CG31674* | -0.55 | 0.68 | 8.18E-03 | uncharacterized protein |
| FBgn0029172 | *Fad2* | -0.79 | 0.58 | 1.60E-05 | Fad2 |
| FBgn0015040 | *Cyp9c1* * | -0.79 | 0.58 | 7.31E-06 | Cytochrome P450-9c1 |
| FBgn0034756 | *Cyp6d2* * | -0.80 | 0.58 | 5.96E-09 | Cyp6d2 |
| FBgn0001112 | *Gld* | -0.95 | 0.52 | 1.43E-06 | Glucose dehydrogenase |
| FBgn0031925 | *Cyp4d21* * | -0.95 | 0.52 | 8.28E-06 | Cyp4d21 |
| FBgn0015714 | *Cyp6a17* * | -1.04 | 0.49 | 2.76E-15 | Cytochrome P450-6a17 |
| FBgn0033395 | *Cyp4p2* * | -2.68 | 0.16 | 3.51E-48 | Cyp4p2 |

\* indicates genes that belong to GO:0020037, GO:0046906, GO:0005506 and GO:0016705.

Among the 220 genes differentially regulated in *para^Shu^* in the absence or presence of *GstS1^M26^* (*para^Shu^*/+; +/+ vs. *para^Shu^*/+; *GstS1^M26^*/+), 25 were up-regulated and 12 were down-regulated (cutoff: fold change >2; Table 7). The gene for which the fold-change was greatest in *para^Shu^* plus *GstS1^M26^* flies was a member of the cytochrome P450 family, *Cyp4p2*; it was down-regulated 6.4-fold in the presence of *GstS1^M26^*, with *P_adj_*=3.5 x 10^-48^. Notably, three of the top 20 genes with the greatest fold expression changes were members of this family (*Cyp4p2*, *Cyp6a8*, *Cyp6a2*).

**Table 7.** Genes most differentially expressed in *para^Shu^*/+; *GstS1^M26^*/+ compared with *para^Shu^*/+; +/+

| Flybase ID | Gene symbol | Fold change (log2) | Fold change | *P*<sub>adj</sub> | Gene product |
| --- | --- | --- | --- | --- | --- |
| FBgn0085732 | *CR40190* | 2.12 | 4.36 | 1.32E-29 | pseudo |
| FBgn0033954 | *CG12860* | 2.02 | 4.05 | 1.42E-26 | uncharacterized protein |
| FBgn0039752 | *CG15530* | 1.95 | 3.86 | 7.79E-30 | uncharacterized protein |
| FBgn0037850 | *CG14695* | 1.90 | 3.73 | 1.32E-29 | uncharacterized protein |
| FBgn0033748 | *vis* | 1.90 | 3.72 | 1.91E-23 | vismay |
| FBgn0266084 | *Fhos* | 1.85 | 3.60 | 6.52E-32 | Formin homology 2 domain containing |
| FBgn0040104 | *lectin-24A* | 1.76 | 3.38 | 5.28E-20 | lectin-24A |
| FBgn0013772 | *Cyp6a8* | 1.69 | 3.22 | 9.38E-20 | Cytochrome P450-6a8 |
| FBgn0031935 | *CG13793* | 1.63 | 3.09 | 1.28E-22 | uncharacterized protein |
| FBgn0000473 | *Cyp6a2* | 1.51 | 2.84 | 6.10E-15 | Cytochrome P450-6a2 |
| FBgn0033926 | *Arc1* | 1.42 | 2.67 | 4.34E-25 | Activity-regulated cytoskeleton associated protein 1 |
| FBgn0085452 | *CG34423* | 1.37 | 2.58 | 1.77E-12 | uncharacterized protein |
| FBgn0259896 | *NimC1* | 1.35 | 2.54 | 1.51E-13 | Nimrod C1 |
| FBgn0261055 | *Sfp26Ad* | 1.28 | 2.43 | 8.50E-11 | Seminal fluid protein 26Ad |
| FBgn0003082 | *phr* | 1.27 | 2.41 | 1.06E-23 | photorepair |
| FBgn0003961 | *Uro* | 1.19 | 2.28 | 1.17E-14 | Urate oxidase |
| FBgn0013308 | *Odc2* | 1.16 | 2.23 | 1.61E-08 | Ornithine decarboxylase 2 |
| FBgn0004426 | *LysC* | 1.12 | 2.18 | 5.61E-08 | Lysozyme C |
| FBgn0052198 | *CG32198* | 1.12 | 2.17 | 3.25E-08 | uncharacterized protein |
| FBgn0041337 | *Cyp309a2* | 1.11 | 2.16 | 1.07E-07 | Cyp309a2 |
| FBgn0053511 | *CG33511* | 1.10 | 2.14 | 1.07E-07 | uncharacterized protein |
| FBgn0034783 | *CG9825* | 1.09 | 2.12 | 1.95E-07 | uncharacterized protein |
| FBgn0032210 | *CYLD* | 1.05 | 2.07 | 7.58E-11 | Cylindromatosis |
| FBgn0033978 | *Cyp6a23* | 1.05 | 2.07 | 7.16E-15 | Cyp6a23 |
| FBgn0004425 | *LysB* | 1.03 | 2.04 | 1.13E-06 | Lysozyme B |
| FBgn0004782 | *Ccp84Ab* | -1.01 | 0.50 | 9.43E-07 | Ccp84Ab |
| FBgn0034715 | *Oatp58Db* | -1.01 | 0.50 | 1.05E-11 | Organic anion<br>transporting polypeptide<br>58Db |
| FBgn0037292 | *plh* | -1.02 | 0.49 | 7.84E-12 | pasang lhamu |
| FBgn0015714 | *Cyp6a17* | -1.04 | 0.49 | 2.76E-15 | Cytochrome P450-6a17 |
| FBgn0030815 | *CG8945* | -1.09 | 0.47 | 3.08E-08 | uncharacterized protein |
| FBgn0004783 | *Ccp84Aa* | -1.10 | 0.47 | 1.11E-07 | Ccp84Aa |
| FBgn0250825 | *CG34241* | -1.21 | 0.43 | 3.92E-10 | uncharacterized protein |
| FBgn0034356 | *Pepck2* | -1.29 | 0.41 | 8.57E-12 | Phosphoenolpyruvate<br>carboxykinase 2 |
| FBgn0031533 | *CG2772* | -1.45 | 0.37 | 4.15E-29 | uncharacterized protein |
| FBgn0031741 | *CG11034* | -1.54 | 0.35 | 1.23E-18 | uncharacterized protein |
| FBgn0260874 | *Ir76a* | -1.68 | 0.31 | 7.94E-19 | Ionotropic receptor 76a |
| FBgn0033395 | *CR40190* | -2.68 | 0.16 | 3.51E-48 | Cyp4p2 |

Listed are genes differentially expressed in *para*<sup>Shu/+</sup> ; *GstS*<sup>1M26/+</sup> compared with *para*<sup>Shu/+</sup> ; +/+ with Fold change > 2 and $P_{adj} < 0.01$ .

## DISCUSSION

In the present study, we performed an unbiased forward genetic screen to identify genes that have a significant impact on the phenotypes associated with *para^Shu^*, a gain-of-function variant of the *Drosophila* Na_v_ channel gene. Our key finding was that a 50% reduction of GstS1 function resulted in strong suppression of *para^Shu^* phenotypes. Glutathione S-transferases (GSTs) are phase II metabolic enzymes that are primarily involved in conjugation of the reduced form of glutathione to endogenous and xenobiotic electrophiles for detoxification (Hayes *et al*. 2005; Allocati *et al*. 2018). Reduced GST function is generally considered damaging to organisms because it is expected to lead to an accumulation of harmful electrophilic compounds in the cell and thereby disturb critical cellular processes. In fact, a previous study showed that loss of *GstS1* function enhanced the loss of dopaminergic neurons in a *parkin* mutant, a *Drosophila* model of Parkinson’s disease and conversely, overexpression of *GstS1* in the same dopaminergic neurons suppressed dopaminergic neurodegeneration in such mutants (Whitworth *et al*. 2005). Parkin has ubiquitin-protein ligase activity (Imai *et al*. 2000; Shimura *et al*. 2000; Zhang *et al*. 2000) and the accumulation of toxic Parkin substrates likely contributes to the degeneration of dopaminergic neurons in Parkinson’s patients and animal models (Whitworth *et al*. 2005). These results are consistent with the idea that GstS1 plays a role in the detoxification of oxidatively damaged products to maintain healthy cellular environments. In this regard, it seems counterintuitive that loss of *GstS1* function reduces, rather than increases, the severity of *para^Shu^* phenotypes.

GstS1 is unique among *Drosophila* GSTs in several respects. A previous study, based on multiple alignments of GST sequences, had revealed that GstS1 is the sole member of the *Drosophila* sigma class of GST (Agianian *et al*. 2003). Unlike other GSTs, GstS1 has low catalytic activity for typical GST substrates, such as 1-chloro-2,4-dinitrobenzol (CDNB), 1,2-dichloro-4-nitrobenzene (DCNB), and ethacrynic acid (EA). Instead, it efficiently catalyzes the conjugation of glutathione to 4-hydroxynonenal (4-HNE), an unsaturated carbonyl compound derived via lipid peroxidation (Singh *et al*. 2001; Agianian *et al*. 2003). The crystal structure of

GstS1 indicates that its active-site topography is suitable for the binding of amphipolar lipid peroxidation products such as 4-HNE (Agianian *et al*. 2003), consistent with the above-mentioned substrate specificity. 4-HNE is the most abundant 4-hydroxyalkenal formed in cells and contributes to the deleterious effects of oxidative stress. It has been implicated in the pathogenesis and progression of human diseases such as cancer, Alzheimer’s disease, diabetes, and cardiovascular disease (Shoeb *et al*. 2014; Csala *et al*. 2015). However, 4-HNE also functions as a signaling molecule and has concentration-dependent effects on various cellular processes including differentiation, growth and apoptosis (Zhang and Forman 2017). GstS1 plays a major role in controlling the intracellular 4-HNE concentration to balance its beneficial and damaging effects; one study estimated that it is responsible for ∼70% of the total capacity to conjugate 4-HNE with glutathione in adult *Drosophila* (Singh *et al*. 2001). It is thus possible that in *para^Shu^* flies the reduction of GstS1 activity enhances the strength of 4-HNE-dependent signaling, leading to changes in neural development and/or function that compensate for the defect caused by the *para^Shu^* mutation.

Notably, GSTs are not limited to conjugating glutathione to potentially harmful substrates for their clearance, and it is possible that another such function accounts for our observations. Specifically, some GSTs catalyze the synthesis of physiologically important compounds. With respect to its primary amino acid sequence, *Drosophila* GstS1 is more similar to the vertebrate hematopoietic prostaglandin D2 synthases (HPGDSs) than to other *Drosophila* GSTs (Agianian *et al*. 2003). Indeed, the sequence identity/similarity between *Drosophila* GstS1 and human HPGDS are 37%/59%, respectively. The *Drosophila* Integrative Ortholog Prediction Tool (DIOPT; Http://www.flyrnai.org/diopt) (Hu *et al*. 2011), as well as a recent and extensive bioinformatics analysis (Scarpati et al. 2019), classified GstS1 as a fly ortholog of HPGDS, a sigma-class member of the GST family that catalyzes the isomerization of prostaglandin H_2_ (PGH_2_) to prostaglandin D_2_ (PGD_2_). Mammalian HPGDS is a critical regulator of inflammation and the innate immune response (Rajakariar *et al*. 2007; Joo and Sadikot 2012). In light of this observation, findings implicating GstS1 in the development and function of the innate immune system in insects are of interest. For example, in a lepidopteran *Spodoptera exigua*, the ortholog of *Drosophila* GstS1, SePGDS, was identified as PGD_2_ synthase, with the addition of PGD_2_, but not its precursor (arachidonic acid), rescuing immunosuppression in larvae in response to SePGDS knockdown (Sajjadian *et al*. 2019). Consistent with this finding, previous studies in *Drosophila* had revealed that overexpression of *GstS1* in hemocytes (the insect blood cells responsible for cellular immunity) leads to increases in the number of larval hemocytes (Stofanko *et al*. 2008) and that GstS1 in hemocytes is increased ∼10-fold at the onset of metamorphosis (Regan *et al*. 2013). These results strongly support a significant role for GstS1 in the insect innate immune system. In addition, we previously found that genes involved in innate immune responses were up-reg ulated in the adult head of *para^Shu^* mutants (Kaas *et al*. 2016), suggesting that the neuronal hyperexcitability induced by gain-of-function *para^Shu^* Na_v_ channels might lead to activation of the innate immune system. In light of these observations and our current findings it is possibile that the reason that loss of GstS1 function reduces the severity of *para^Shu^* phenotypes is that it suppresses the innate immune response through hemocytes and prostaglandin-like bioactive lipids.

Another connection to the innate immune system is the discovery, based on our transcriptome analysis, that CYP genes are over-represented among the genes that are differentially expressed in the *para^Shu^* with a *GstS1* mutation (Table 5). CYP enzymes are involved in the oxygenation of a wide range of compounds, including eicosanoids such as prostaglandins. In mammals, activation of the innate immune response alters CYP expression and eicosanoid metabolism in an isoform-, tissue-, and time-dependent manner (Theken *et al*. 2011). *GstS1* loss of function may affect *para^Shu^* phenotypes by changing the activities of CYP enzymes. Further studies are required to elucidate whether and how CYP genes, as well as the genes involved in innate immune response and bioactive lipid signaling, contribute to *GstS1*-mediated modulation of *para^Shu^* phenotypes.

To obtain insight into functional significance of changes in gene expression, we classified differentially expressed genes. For the 89 genes that were up-regulated by *para^Shu^* (*para^Shu^*/+ vs. +/+), it is notable that 13 were down-regulated when *GstS1^M26^* was also introduced (*para^Shu^*/+ vs. *para^Shu^*/+; *GstS1^M26^*/+) and that all of the GO categories associated (*P_adj_*<0.05) with this group of genes were related to the chitin-based cuticle (Table 3A). On the other hand, among the 40 genes down-regulated by *para^Shu^*, only 2 (*CG5966* and *CG5770*) were up-regulated by *GstS1^M26^*. Although *CG5770* is an uncharacterized gene, *CG5966* encodes proteins that are highly expressed in the larval and adult fat bodies and predicted to be involved in lipid catabolism. A human *CG5966* homolog encodes pancreatic lipase, which hydrolyzes triglycerides in the small intestine and is essential for the efficient digestion of dietary fat (Davis *et al*. 1991). Notably, changes in the expression of these cuticle-associated and fat metabolism-associated sets of genes appear to correlate with the phenotypic severity of *para^Shu^* in that a change in the phenotype or gene expression induced by *para^Shu^* is reversed by *GstS1^M26^*. It is possible that changes in the expression of these genes is causative and contributes to the severity of *para^Shu^* phenotypes. Alternatively, these changes in gene expression could be a consequence of phenotypic changes caused by other factors. Further functional analysis is required to determine the significance of these genes in controlling *para^Shu^* phenotypes.

In contrast to the expression of the above-mentioned genes, that of 24 genes was changed in the same direction by *para^Shu^* and *GstS1^M26^*. Among these, 17 were up-regulated and 7 were down-regulated. No GO category was identified for any of the gene sets with *P_adj_*<0.05. Interestingly, *GstS1* itself is one of the genes whose expression is down-regulated by both *para^Shu^* and *GstS1^M26^*. The observed reduction in levels of *GstS1* expression in the *GstS1^M26^* mutant is consistent with it being a deletion allele. However, its down-regulation in *para^Shu^* mutants was unexpected. One possible explanation for this finding is that homeostatic regulation at the level of gene expression counteracts the defects caused by hyperexcitability. It will be important to elucidate the mechanisms by which a gain-of-function mutation in a Na_v_-channel gene leads to down-regulation of the expression of its modifier gene and to reduction of the severity of the phenotype.

A previous genetic screen that was similar to ours revealed that loss of the function of *gilgamesh (gish)* reduces the severity of the seizure phenotypes of *para^bss^* mutant. *gish* encodes the *Drosophila* ortholog of casein kinase CK1γ3, a member of the CK1 family of serine-threonine kinases (Howlett *et al*. 2013). Another modifier of seizure activity was discovered by Lin et al. (2017); this group identified *pumilio* (*pum*) based on transcriptome analyses of *Drosophila* seizure models, with *pum* significantly down-regulated in both the genetic (*para^bss^*) and pharmacological (picrotoxin-induced) models. It was shown that pan-neuronal overexpression of *pum* is sufficient to dramatically reduce seizure severity in *para^bss^* as well as other seizure-prone *Drosophila* mutants, *easily shocked* (*eas*) and *slamdance* (*sda*) (Lin *et al*. 2017). *pum* encodes RNA binding proteins that act as homeostatic regulators of action potential firing, partly by regulating the translation of *para* transcripts (Lin *et al*. 2017). In addition, we recently discovered that the seizure phenotypes of *para^Shu^* and other seizure-prone fly mutants are significantly suppressed when the flies are fed a diet supplemented with milk whey (Kasuya *et al*. 2019). It remains unclear how these genetic and environmental factors interact with one another in complex regulatory networks and how they modify the neurological phenotypes of mutants. A mechanistic understanding of such functional interactions is expected to reveal the molecular and cellular processes that are critical for the manifestation of hyperexcitable phenotypes in *Drosophila* mutants, and to provide useful insights into the corresponding processes in vertebrate animals, including humans.

## Acknowledgements

We thank Mr. Ryan Jewell and Mr. Pei-Jen Wang (Department of Medical Research, Tungs’ Taichung MetroHarbor Hospital, Taichung City, Taiwan 43503, ROC) for their technical assistance.

