## Supplemental Table 1 for "Reduced function of the glutathione S-transferase S1 suppresses behavioral hyperexcitability in *Drosophila* expressing a mutant voltage-gated sodium channel"

| **2nd chromosome** | | | |
| --- | --- | --- | --- |
| **Df Name** | **deletion location** | **CI** | **N** |
| *T(1;2)l-v75, y[1]/FM6* | 19-20;41 | 0.00 | 7 |
| *Df(2L)net-PMF* | 21A1;21B7-21B8 | 0.04 | 14 |
| *Df(2L)net-PMF* | 21A1;21B7-8 | 0.00 | 6 |
| *Df(2L)BSC4* | 21B7-C1;21C2-3 | 0.04 | 15 |
| *Df(2L)BSC16* | 21C3-4;21C6-8 | 0.13 | 8 |
| *Df(2L)ast2* | 21D1-2;22B2-3 | 0.09 | 11 |
| *Df(2L)dp-79b* | 22A2-3;22D5-E1 | 0.29 | 14 |
| *Df(2L)BSC37* | 22D2-3;22F1-2 | 0.09 | 11 |
| *Df(2L)dpp[d14]* | 22E4-F2;22F3-23A1 | 0.00 | 4 |
| *Df(2L)C144* | 22F4-23A1;23C2-4 | 0.02 | 9 |
| *Df(2L)JS17* | 23C1-2;23E1-2 | 0.38 | 12 |
| *Df(2L)BSC28* | 23C5-D1;23E2 | 0.73 | 12 |
| *Df(2L)BSC31* | 23E5;23F4-5 | 0.22 | 12 |
| *Df(2L)drm-P2* | 23F3-4;24A1-2 | 0.00 | 4 |
| *Df(2L)ed1* | 24A2;24D4 | 0.03 | 7 |
| *Df(2L)sc19-8* | 24C2-8;25C8-9 | 0.40 | 9 |
| *Df(2L)cl-h3* | 25D2-4;26B2-5 | 0.00 | 4 |
| *Df(2L)E110* | 25F3-26A1;26D3-11 | 0.30 | 4 |
| *Df(2L)BSC5* | 26B1-2;26D1-2 | 0.00 | 17 |
| *Df(2L)BSC7* | 26D10-E1;27C1 | 0.13 | 8 |
| *Df(2L)BSC6* | 26D3-E1;26F4-7 | 0.11 | 7 |
| *Df(2L)spd[j2]* | 27C1-2;28A | 0.26 | 19 |
| *Df(2L)Dwee1-W05* | 27C2-3;27C4-5 | 0.33 | 26 |
| *Df(2L)XE-3801* | 27E2;28D1 | 0.00 | 11 |
| *Df(2L)BSC41* | 28A4-B1;28D3-9 | 0.03 | 15 |
| *Df(2L)Trf-C6R31* | 28DE;28DE | 0.20 | 11 |
| *Df(2L)TE29Aa-11* | 28E4-7;29B2-C1 | 0.00 | 4 |
| *Df(2L)N22-14* | 29C1-2;30C8-9 | 0.30 | 6 |
| *Df(2L)BSC17* | 30C3-5;30F1 | 0.13 | 6 |
| *Df(2L)Mdh* | 30D-30F;31F | 0.04 | 5 |
| *Df(2L)BSC50* | 30F4-5;31B1-4 | 0.19 | 23 |
| *Df(2L)J2* | 31B;32A | 0.32 | 5 |
| *Df(2L)BSC32* | 32A1-2;32C5-D1 | 0.43 | 7 |
| *Df(2L)BSC36* | 32D1;32D4-E1 | 0.36 | 16 |
| *Df(2L)FCK-20* | 32D1;32F1-3 | 0.10 | 6 |
| *Df(2L)Prl* | 32F1-3;33F1-2 | 0.03 | 16 |
| *Df(2L)BSC30* | 34A3;34B7-9 | 0.08 | 5 |
| *Df(2L)b87e25* | 34B12-C1;35B10-C1 | 0.00 | 8 |
| *Df(2L)fn2* | 35A3;35B2 | 0.03 | 15 |
| *Df(2L)Exel6035* | 35A3;35B2 | 0.00 | 9 |
| *Df(2L)r10* | 35D1;36A6-7 | 0.00 | 9 |
| *Df(2L)cact-255rv64* | 35F-36A;36D | 0.05 | 11 |
| *Df(2L)C'* | h35;h38L | 0.25 | 13 |
| *Df(2L)TW137* | 36C2-4;37B9-C1 | 0.07 | 9 |
| *Df(2L)pr-A16* | 37B2-12;38D2-5 | 0.56 | 5 |
| *Df(2L)TW161* | 38A6-B1;40A4-B1 | 0.23 | 8 |
| *Df(2R)M41A4* | 41A;41A | 0.00 | 7 |
| *Df(2R)nap9* | 42A1-2;42E6-F1 | 0.00 | 7 |
| *Df(2R)ST1* | 42B3-5;43E15-18 | 0.00 | 10 |
| *In(2R)bw[VDe2L]Cy[R]* | h42-h43;42A2-3 | 0.04 | 10 |
| *Df(2R)H3C1* | 43F;44D3-8 | 0.59 | 19 |
| *Df(2R)H3E1* | 44D1-4;44F12 | 0.22 | 20 |
| *Df(2R)Np5* | 44F10;45D9-E1 | 0.16 | 10 |
| *Df(2R)w45-30n* | 45A6-7;45E2-3 | 0.00 | 9 |
| *Df(2R)BSC29* | 45D3-4;45F2-6 | 0.15 | 4 |
| *Df(2R)B5* | 46A;46C | 0.00 | 8 |
| *Df(2R)en-A* | 47D3;48B2 | 0.00 | 4 |
| *Df(2R)en30* | 48A3-4;48C6-8 | 0.11 | 19 |
| *Df(2R)BSC39* | 48C5-D1;48D5-E1 | 0.04 | 11 |
| *Df(2R)CB21* | 48E;49A | 0.07 | 15 |
| *Df(2R)BSC40* | 48E1-2;48E2-10 | 0.10 | 10 |
| *Df(2R)vg-C* | 49A4-13;49E7-F1 | 0.07 | 12 |
| *Df(2R)CX1* | 49C1-4;50C23-D2 | 0.03 | 14 |
| *Df(2R)BSC18* | 50D1;50D2-7 | 0.04 | 11 |
| *Df(2R)BSC11* | 50E6-F1;51E2-4 | 0.03 | 12 |
| *Df(2R)Jp1* | 51D3-8;52F5-9 | 0.00 | 1 |
| *Df(2R)Jp8* | 52F5-9;52F10-53A1 | 0.07 | 9 |
| *Df(2R)BSC49* | 53D9-E1;54B5-10 | 0.28 | 5 |
| *Df(2R)P803-Delta15* | 53E;53F11 | 0.40 | 18 |
| *Df(2R)ED1* | 53E4;53F8 | 0.05 | 8 |
| *Df(2R)BSC44* | 54B1-2;54B7-10 | 0.08 | 16 |
| *Df(2R)robl-c* | 54B17-C4;54C1-4 | 0.02 | 9 |
| *Df(2R)k10408* | 54C1-4;54C1-4 | 0.00 | 8 |
| *Df(2R)BSC45* | 54C8-D1;54E2-7 | 0.00 | 15 |
| *Df(2R)14H10Y-53* | 54D1-2;54E5-7 | 0.16 | 11 |
| *Df(2R)14H10W-35* | 54E5-7;55B5-7 | 0.20 | 13 |
| *Df(2R)PC4* | 55A;55F | 0.04 | 5 |
| *Df(2R)P34* | 55E2-4;56C1-11 | 0.04 | 11 |
| *Df(2R)BSC26* | 56C4;56D6-10 | 0.00 | 4 |
| *Df(2R)BSC22* | 56D7-E3;56F9-12 | 0.03 | 8 |
| *Df(2R)BSC19* | 56F12-14;57A4 | 0.07 | 6 |
| *Df(2R)017/SM1* | 56F5;56F15 | 0.04 | 15 |
| *Df(2R)AA21* | 56F9-17;57D11-12 | 0.00 | 5 |
| *Df(2R)Egfr5* | 57D2-8;58D1 | 0.15 | 13 |
| *Df(2R)59AD* | 59A1-3;59D1-4 | 0.00 | 5 |
| *Df(2R)vir130* | 59B;59D8-E1 | 0.30 | 14 |
| *Df(2R)Px2* | 60C5-6;60D9-10 | 0.00 | 8 |
| *Df(2R)M60E* | 60E2-3;60E11-12 | 0.20 | 9 |
| *CS* | N/A | 0.04 | 65 |

CI: Climbing Index

N: Number of flies tested

| **3rd chromosome** | | | |
| --- | --- | --- | --- |
| **Df Name** | **deletion location** | **CI** | **N** |
| *Df(3L)emc-E12* | 61A;61D3 | 0.00 | 18 |
| *Df(3L)Ar14-8* | 61C5-8;62A8 | 0.00 | 18 |
| *Df(3L)Aprt-1* | 62A10-B1;62D2-5 | 0.00 | 26 |
| *Df(3L)R-G7* | 62B4-7;62D5-E5 | 0.04 | 17 |
| *Df(3L)BSC23* | 62E8;63B5-6 | 0.00 | 14 |
| *Df(3L)GN34* | 63E6-9;64A8-9 | 0.00 | 20 |
| *Df(3L)GN24* | 63F6-7;64C13-15 | 0.00 | 21 |
| *Df(3L)ZN47* | 64C;65C | 0.18 | 27 |
| *Df(3L)XDI98* | 65A2;65E1 | 0.09 | 16 |
| *Df(3L)BSC27* | 65D4-5;65E4-6 | 0.00 | 16 |
| *Df(3L)BSC33* | 65E10-F1;65F2-6 | 0.00 | 27 |
| *Df(3L)pbl-X1* | 65F3;66B10 | 0.00 | 18 |
| *Df(3L)ZP1* | 66A17-20;66C1-5 | 0.10 | 50 |
| *Df(3L)BSC13* | 66B12-C1;66D2-4 | 0.04 | 15 |
| *Df(3L)66C-G28* | 66B8-9;66C9-10 | 0.00 | 20 |
| *Df(3L)h-i22* | 66D10-11;66E1-2 | 0.00 | 11 |
| *Df(3L)Scf-R6* | 66E1-6;66F1-6 | 0.12 | 31 |
| *Df(3L)BSC35* | 66F1-2;67B2-3 | 0.00 | 54 |
| *Df(3L)AC1* | 67A2;67D11-13 | 0.34 | 29 |
| *Df(3L)BSC14* | 67E3-7;68A2-6 | 0.00 | 15 |
| *Df(3L)vin5* | 68A2-3;69A1-3 | 0.00 | 17 |
| *Df(3L)vin7* | 68C8-11;69B4-5 | 0.02 | 18 |
| *Df(3L)eyg[C1]* | 69A4-5;69D4-6 | 0.00 | 24 |
| *Df(3L)BSC10* | 69D4-5;69F5-7 | 0.06 | 18 |
| *Df(3L)BSC12* | 69F6-70A1;70A1-2 | 0.01 | 21 |
| *In(3LR)C190[L]Ubx[42TR]* | 70A1-2;70C3-4 | 0.00 | 19 |
| *Df(3L)fz-GF3b* | 70C1-2;70D4-5 | 0.14 | 17 |
| *Df(3L)fz-M21* | 70D2-3;71E4-5 | 0.00 | 12 |
| *Df(3L)XG5* | 71C2-3;72B1-C1 | 0.00 | 11 |
| *Df(3L)BSC8* | 74D3-75A1;75B2-5 | 0.00 | 1 |
| *Df(3L)Cat* | 75B8;75F1 | 0.00 | 15 |
| *Df(3L)fz2* | 75F10-11;76A1-5 | 0.00 | 15 |
| *Df(3L)ED4782* | 75F2;76A1 | 0.00 | 11 |
| *Df(3L)XS533* | 76B4;77B | 0.35 | 57 |
| *Df(3L)rdgC-co2* | 77A1;77D1 | 0.05 | 21 |
| *Df(3L)ri-79c* | 77B-C;77F-78A | 0.23 | 59 |
| *Df(3L)ri-XT1* | 77E2-4;78A2-4 | 0.02 | 29 |
| *Df(3L)ME107* | 77F3;78C8-9 | 0.09 | 16 |
| *Df(3L)Pc-2q* | 78C5-6;78E3-79A1 | 0.13 | 11 |
| *Df(3L)ED4978* | 78D5;79A2 | 0.04 | 22 |
| *Df(3L)Ten-m-AL29* | 79C1-3;79E3-8 | 0.01 | 17 |
| *Df(3L)HD1* | 79D3-E1;79F3-6 | 0.00 | 16 |
| *Df(3L)BSC21* | 79E5-F1;80A2-3 | 0.08 | 89 |
| *Df(3R)ME15* | 81F3-6;82F5-7 | 0.42 | 62 |
| *Df(3R)3-4* | 82F3-4;82F10-11 | 0.01 | 17 |
| *Df(3R)e1025-14* | 82F8-10;83A1-3 | 0.15 | 14 |
| *Df(3R)ED5177* | 83B4;83B6 | 0.02 | 22 |
| *Df(3R)BSC47* | 83B7-C1;83C6-D1 | 0.16 | 38 |
| *Df(3R)Tpl10* | 83C1-2;84B1-2 | 0.07 | 17 |
| *Df(3R)WIN11* | 83E1-2;84A5 | 0.01 | 15 |
| *Df(3R)Scr* | 84A1-2;84B1-2 | 0.00 | 15 |
| *Df(3R)Antp17* | 84A5;84D9 | 0.00 | 20 |
| *Df(3R)p-XT103* | 85A2;85C1-2 | 0.01 | 41 |
| *Df(3R)BSC24* | 85C4-9;85D12-14 | 0.49 | 67 |
| *Df(3R)by10* | 85D8-12;85E7-F1 | 0.08 | 17 |
| *Df(3R)BSC38* | 85F1-2;86C7-8 | 0.05 | 15 |
| *Df(3R)M-Kx1* | 86C1;87B1-5 | 0.00 | 18 |
| *Df(3R)T-32* | 86E2-4;87C6-7 | 0.21 | 14 |
| *Df(3R)ry615* | 87B11-13;87E8-11 | 0.43 | 30 |
| *Df(3R)ea* | 88E7-13;89A1 | 0.07 | 15 |
| *Df(3R)sbd105* | 88F9-89A1;89B9-10 | 0.03 | 18 |
| *Df(3R)sbd104* | 89B5;89C2-7 | 0.43 | 15 |
| *Df(3R)Cha7* | 90F1-F4;91F5 | 0.00 | 20 |
| *Df(3R)BSC43* | 92F7-93A1;93B3-6 | 0.05 | 13 |
| *Df(3R)e-N19* | 93B;94 | 0.02 | 27 |
| *Df(3R)e-R1* | 93B6-7;93D2 | 0.00 | 13 |
| *Df(3R)BSC55* | 94D2-10;94E1-6 | 0.12 | 20 |
| *Df(3R)BSC56* | 94E1-2;94F1-2 | 0.26 | 21 |
| *Df(3R)Exel9012* | 94E9;94E13 | 0.01 | 17 |
| *Df(3R)BSC137* | 95A2-4;95A8-B1 | 0.02 | 20 |
| *Df(3R)mbc-30* | 95A5-7;95C10-11 | 0.00 | 26 |
| *Df(3R)mbc-R1* | 95A5-7;95D6-11 | 0.07 | 18 |
| *Df(3R)crb-F89-4* | 95D7-D11;95F15 | 0.00 | 19 |
| *Df(3R)crb87-5* | 95F7;96A17-18 | 0.00 | 17 |
| *Df(3R)slo8* | 96A2-7;96D2-4 | 0.00 | 2 |
| *Df(3R)BSC140* | 96F1;96F10 | 0.05 | 26 |
| *Df(3R)Espl3* | 96F1;97B1 | 0.00 | 6 |
| *Df(3R)Tl-P* | 97A;98A1-2 | 0.00 | 15 |
| *Df(3R)D605* | 97E3;98A5 | 0.02 | 48 |
| *Df(3R)BSC42* | 98B1-2;98B3-5 | 0.00 | 15 |
| *Df(3R)3450* | 98E3;99A6-8 | 0.00 | 20 |
| *Df(3R)Dr-rv1* | 99A1-2;99B6-11 | 0.02 | 23 |
| *Df(3R)B81* | 99D3;3Rt | 0.13 | 31 |
| *CS* | N/A | 0.04 | 65 |

CI: Climbing Index

N: Number of flies tested
